# Inference of the demographic histories and selective effects of human gut commensal microbiota over the course of human history

**DOI:** 10.1101/2023.11.09.566454

**Authors:** Jonathan C. Mah, Kirk E. Lohmueller, Nandita Garud

**Author notes:** Co-correspondence to and.

## Abstract

Despite the importance of gut commensal microbiota to human health, there is little knowledge about their evolutionary histories, including their population demographic histories and their distributions of fitness effects (DFE) of new mutations. Here, we infer the demographic histories and DFEs of 27 of the most highly prevalent and abundant commensal gut microbial species in North Americans over timescales exceeding human generations using a collection of lineages inferred from a panel of healthy hosts. We find overall reductions in genetic variation among commensal gut microbes sampled from a Western population relative to an African rural population. Additionally, some species in North American microbiomes display contractions in population size and others expansions, potentially occurring at several key historical moments in human history. DFEs across species vary from highly to mildly deleterious, with accessory genes experiencing more drift compared to core genes. Within genera, DFEs tend to be more congruent, reflective of underlying phylogenetic relationships. Taken together, these findings suggest that human commensal gut microbes have distinct evolutionary histories, possibly reflecting the unique roles of individual members of the microbiome.

## INTRODUCTION

Diversity in the human gut microbiome varies significantly across geographic regions around the world. Notably, rural African microbiomes display greater levels of diversity than urban Western microbiomes at multiple levels including at the species (Sonnenburg et al. 2016, Nature), gene (Tett et al. 2019, Cell Host Microbe), and nucleotide (Nayfach and Pollard 2015, bioRxiv; Tett et al. 2019, Cell Host Microbe) levels. A number of demographic and selective forces may be responsible for these differences in diversity, including urbanization (Obregon-Tito et al. 2015, Nature Comm), antibiotic usage (Sonnenberg and Sonnenberg 2019, Science; Clemente et al. 2015, Science Advances), shifts in diet (Blaser et al. 2018, Cell; Schnorr et al. 2014 Nature Comm; De Filippo et al. 2010, PNAS), and human migration (Yatsunenko et al. 2012, Nature). For example, reductions in dietary fiber in mice lead to irreversible reductions in genetic diversity in the gut microbiome and are thought to be a contributing factor to the depleted diversity of rural agrarian populations compared to hunter-gatherer populations (Sonnenburg et al. 2016, Nature). Additionally, some microbes such as *Helicobacter pylori* (Falush et al. 2003, Science) and *Prevotella copri* (Tett et al. 2019, Cell Host Microbe) display phylogenetic patterns mirroring human migration events around the world, suggesting that human demographic events themselves may have shaped bacterial genomic variation.

To date, there has not been a systematic inference of the demographic histories and distributions of fitness effects (DFEs) of new mutations of commensal gut microbiota found across hosts, nor has there been any investigation into how these quantities might vary across species and genera. Such inferences are important for a number of reasons. Demographic inference provides an understanding of pre-historical events, such as continental migrations as well as population contractions and expansions. Additionally, demographic inference enables the ability to detect genomic regions subject to selective forces by providing an expectation of genetic diversity under neutral conditions. Inference of the DFE, which quantifies the proportion of new mutations that are deleterious, neutral, or beneficial, is necessary for addressing fundamental questions such as the fate of colonizing lineages within a human host and the evolutionary capacity of a population to respond to novel selection pressures (Chevin et al. 2010, PLoS Biology; Dapa et al. 2023, Current Opinion). Finally, a better understanding of how and why microbiomes across diverse human populations differ might aid precision health purposes, especially if strains within species interact differently with their hosts across populations (Yatsunenko et al. 2012, Nature).

A key population genetics approach for the inference of demographic histories and DFEs among eukaryotes leverages a summary statistic known as the site frequency spectrum or SFS (**Figure 2**) (Wakeley and Hey 1997, Genetics; Nielsen 2000, Genetics; Gutenkunst et al. 2009, PLoS Genetics). The SFS describes the distribution of minor allele frequencies from a given sample of DNA sequences and is highly sensitive to the impacts of demography and selection (Nielsen 2000, Genetics). For example, relative to the SFS of a population in demographic equilibrium, a population undergoing a demographic expansion and/or purifying selection would harbor a higher proportion of rare variants, i.e., SNPs present in a small number of individuals relative to the overall sample size. Conversely, a population undergoing a demographic contraction and/or balancing selection would harbor a higher proportion of common variants.

While the SFS has been extensively used to infer demography and selective effects in eukaryotes (Beichman et al 2018, Annual Reviews; Gutenkunst et al 2009, PLoS Genetics; Boyko et al 2008, PLoS Genetics; Eyre-Walker et al. 2006, Genetics), it has been under-utilized for bacteria (Pepperell et al 2013, PLoS Pathogens; Cornejo et al 2013, MBE). Yet, with this approach, the cavity-causing oral bacterium *Streptococcus mutans* was inferred to have undergone a demographic expansion, coincident with the onset of agricultural development ∼ 10,000 – 20,000 years ago (Cornejo et al. 2013, MBE), potentially due to an expanded niche via new available resources from the shift in diet. Additionally, mutations in this species, like those in many other bacterial species, were inferred to experience extensive purifying selection using the SFS. Application of SFS-based analyses to human gut commensal bacteria found across a panel of hosts may similarly yield insights into demographic events and DFEs of new mutations on time scales exceeding human generations.

One important assumption of applying an SFS-based analysis is quasi-independence between sites, i.e., free recombination between sites (Bustamante et al. 2001, Genetics). While bacteria reproduce asexually, most commensal gut bacteria experience extensive recombination (Vos and Didelot 2009, ISME; Liu and Good 2023, bioRxiv; Garud and Good et al. 2019, PLoS Biology; Lin and Kussel 2019, Nature Methods; Smith et al. 1993, PNAS). This recombination plays a critical role in shuffling genetic material and breaking up correlations between loci in the genome on long timescales exceeding within-host evolution. The effects of recombination on bacterial genomic diversity is evident in the strong decay in linkage disequilibrium (LD) with genomic distance in samples of lineages from human hosts around the world (Garud and Good et al. 2019, PLoS Biology; Liu and Good 2023, bioRxiv; Sakoparnig et al. 2021, Elife; Lin and Kussel 2019, Nature Methods), as well as phylogenetic inconsistencies at the SNP and gene level with trees built from genome-wide data (Sakoparnig et al. 2021, Elife). It has been estimated that recombination contributes on the order of 10 times more to variation than mutations to gut bacteria, indicating that recombination overwrites any semblance of clonality of circulating lineages (Liu and Good 2023, bioRxiv). Given the relatively high rates of recombination in gut microbiota, we propose to use the SFS to gain insights into the evolutionary history of commensal bacteria.

In this study, we perform an SFS-based inference of the demographic history and fitness effects acting on 27 prevalent commensal gut species using a panel of healthy hosts from North America. Concretely, we examine the evolutionary dynamics operating across hosts on timescales exceeding a human lifetime by building SFSs composed of a single strain per individual microbiome for each species. We infer that human gut commensal bacteria have experienced a range of demographic histories, including contractions and expansions in effective population size over the past ∼1,000 – 100,000 years of human history. Additionally, we leverage the SFS to infer the DFE, finding that bacterial DFEs are more congruent within versus between genera, indicative of underlying phylogenetic relationships. Finally, we find that accessory genes experience more drift compared to core genes. Taken together, these findings suggest that common commensal gut microbiota have distinct evolutionary histories.

## RESULTS

### Data

To infer the demographic histories and distribution of fitness effects of new mutations of gut commensal bacteria found in Western urbanized populations, we analyzed shotgun metagenomic data from a panel of 250 healthy hosts from North America from the Human Microbiome Project (HMP) (**Table S1**). Additionally, we analyzed shotgun metagenomic data from 121 healthy hosts from a rural African cohort from Northeastern Madagascar (Pasolli et al. 2019, Cell) (**Table S2**). To identify single nucleotide variants (SNVs) and gene copy number variants (CNVs) from these metagenomic samples, reads were mapped to reference genomes using a standard pipeline (Nayfach et al. 2016, Genome Research; Methods).

### Depletion of nucleotide diversity in North American gut microbiomes relative to rural African gut microbiomes

To gain an initial understanding into the demographic history of commensal gut microbiota from healthy North American individuals, we compare their nucleotide diversity (π) with that of commensal gut microbiota from healthy rural African individuals (Methods). This analysis is motivated by previous work showing that microbiota from African individuals were a potential ancestral population for gut microbiota in non-African individuals (Suzuki et al. 2022, Science; Moeller et al. 2016, Science), including species such as *Prevotella copri* (Tett et al. 2019, Cell Host Microbe), and *Helicobacter pylori* (Falush et al. 2003, Science). Specifically, to gain insight into changes in effective population sizes in North American gut microbiota compared to African gut microbiota, we analyze within-host and between-host nucleotide diversity (π) for all prevalent species found in either cohort.

Mean within-host nucleotide diversity in North Americans is lower on average compared to within African individuals (**Figure 1**). Although most species that are prevalent in North American hosts (n=81 species present in 5 or more individuals) do not overlap with species that are prevalent in African hosts (n=128 species present in 5 or more individuals), of the 13 species present in at least 5 North American and 5 African gut microbiomes, 9 have significantly decreased within-host nucleotide diversity in North American gut microbiomes than in African gut microbiomes (P-value =< 0.05; two-sided Wilcoxon rank-sum test) (**Figure 1A**).

**Figure 1:**
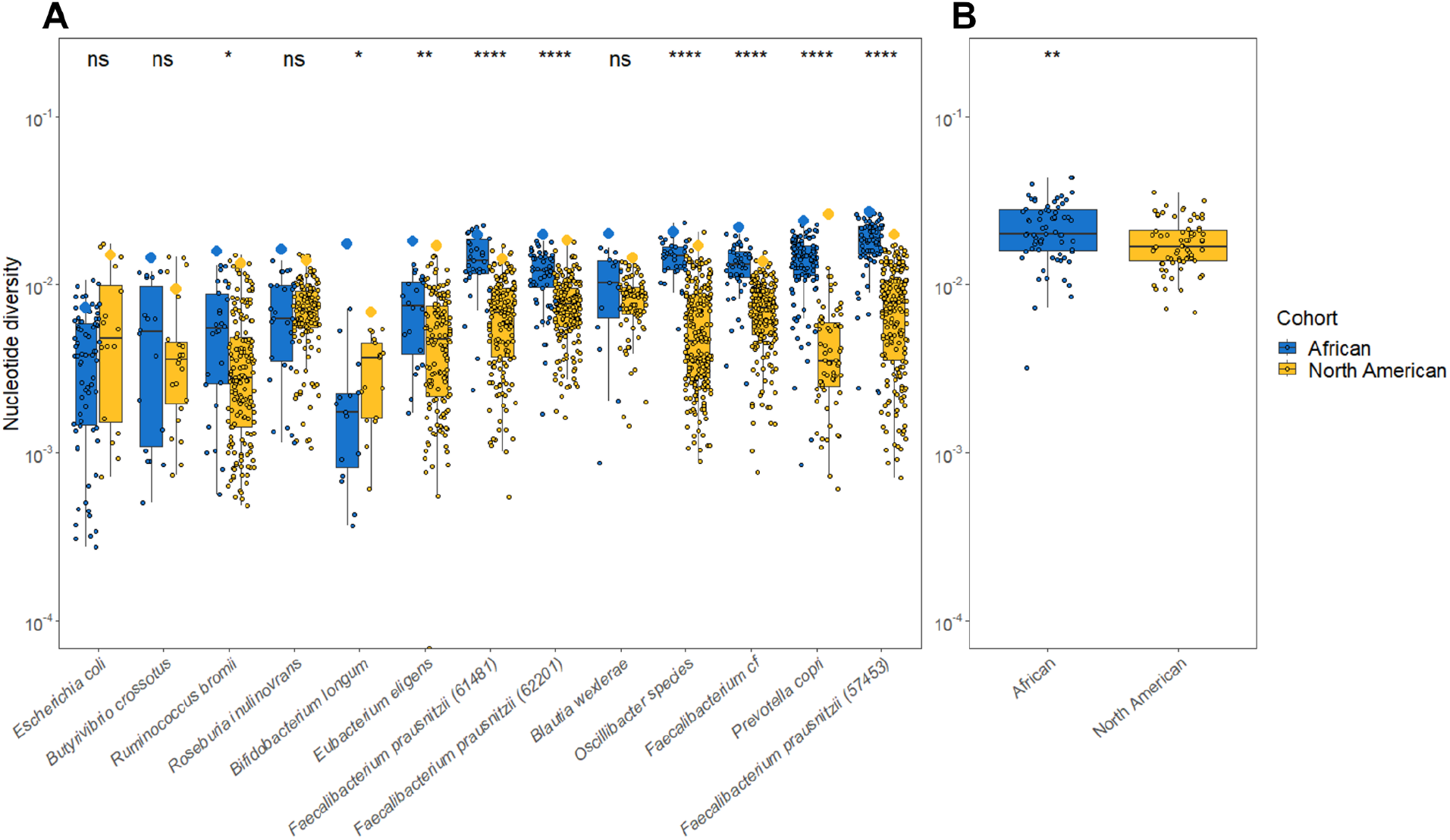
Nucleotide diversity within and between hosts in rural African and North American gut microbiomes. **(A)** Each dot represents within-host nucleotide diversity of a single host; larger diamonds represent mean between-host nucleotide diversity across all pairs of hosts. Nine species have a significant difference in the distribution of within-host nucleotide diversity between North Americans and Africans (ns: not significant, *: P-value =< 0.05, **: P-value =< 0.01, ***: P-value =< 0.001; two-sided Wilcoxon rank sum test). Diversity distributions are shown only for those species present in at least 5 African and 5 North American hosts. **(B)** Distribution of between-host nucleotide diversity of prevalent species found in at least 5 African or 5 North American hosts. Each dot represents mean between-host nucleotide diversity for a given species. Mean between-host nucleotide diversity in the African cohort is significantly higher than that of the North American cohort (P-value = 3.524 x 10^-3^; two-sided Wilcoxon rank sum test).

This overall lower mean within-host nucleotide diversity in North Americans likely reflects fewer strains per species on average colonizing a typical North American host compared to a typical African host. Given conservatively high estimates of mutation rates, generation times, and number of years within a host, it has been estimated that a single lineage cannot generate more than 10^-3^ /bp nucleotide diversity in a host’s lifetime (Garud and Good et al. 2019, PLoS Biology). Hence, higher levels of within-host nucleotide diversity (e.g. on the order of 10^-2^/bp) are unlikely to be explained by the evolution of single lineages within a host and are instead better accounted for by the presence of multiple co-colonizing strains. However, hosts with species displaying lower levels of within-host diversity are likely colonized by a single lineage of that species.

Additionally, mean between-host diversity computed between pairs of individuals (Methods) is also significantly depleted in North American gut microbiomes relative to African gut microbiomes for these same species (P-value = 2.66 x 10^-2^, paired two-sided Wilcoxon signed rank test). More generally, for all species found in either cohort in at least 5 individuals, irrespective of their overlap, mean between-host nucleotide diversity is significantly depleted in North American gut microbiomes relative to African gut microbiomes (P-value = 3.52 x 10^-3^, unpaired two-sided Wilcoxon rank-sum test, **Figure 1B**).

Together these findings suggest that, on average, per-species nucleotide diversity is depleted both within and between hosts in North American gut microbiota relative to African gut microbiota. This depletion of diversity within North American hosts suggests that they, on average, harbor fewer strains per species as compared to African hosts. Additionally, the reduced diversity observed between hosts in North America versus Africa indicates an overall lower effective population size in North American gut microbiomes. These findings are consistent with previous work showing a general reduction in microbiome diversity in North Americans compared to rural Africans at the species level (Yatsunenko et al. 2012, Nature; Moeller et al. 2014, PNAS; Moeller et al. 2016, Science; Sonnenberg et al. 2016, Nature; Schnorr et al. 2014, Nature Communications; Blaser et al. 2018, Cell) as well as at the nucleotide level (Nayfach and Pollard 2015, BioRxiv; Tett et al. 2019, Cell Host Microbe; Karcher et al. 2020, Genome Biology), and indicate that evolutionary processes, such as demographic changes or purifying selection, may be responsible for differences in diversity levels in North American microbiomes relative to rural African microbiomes.

### Gut microbiota experience a range of demographic changes on time scales spanning human history

#### Building SFSs from metagenomic data from a panel of hosts

We next investigated the demographic histories of the 27 most prevalent and abundant commensal gut microbiota found in North Americans (Figure 2). Specifically, we investigated long-term demographic changes occurring on timescales exceeding a human lifetime as opposed to demographic changes occurring within a host’s lifetime. As such, we analyzed SFSs composed of lineages from different hosts as opposed to within-host SFSs (**Figure 2**).

**Figure 2:**
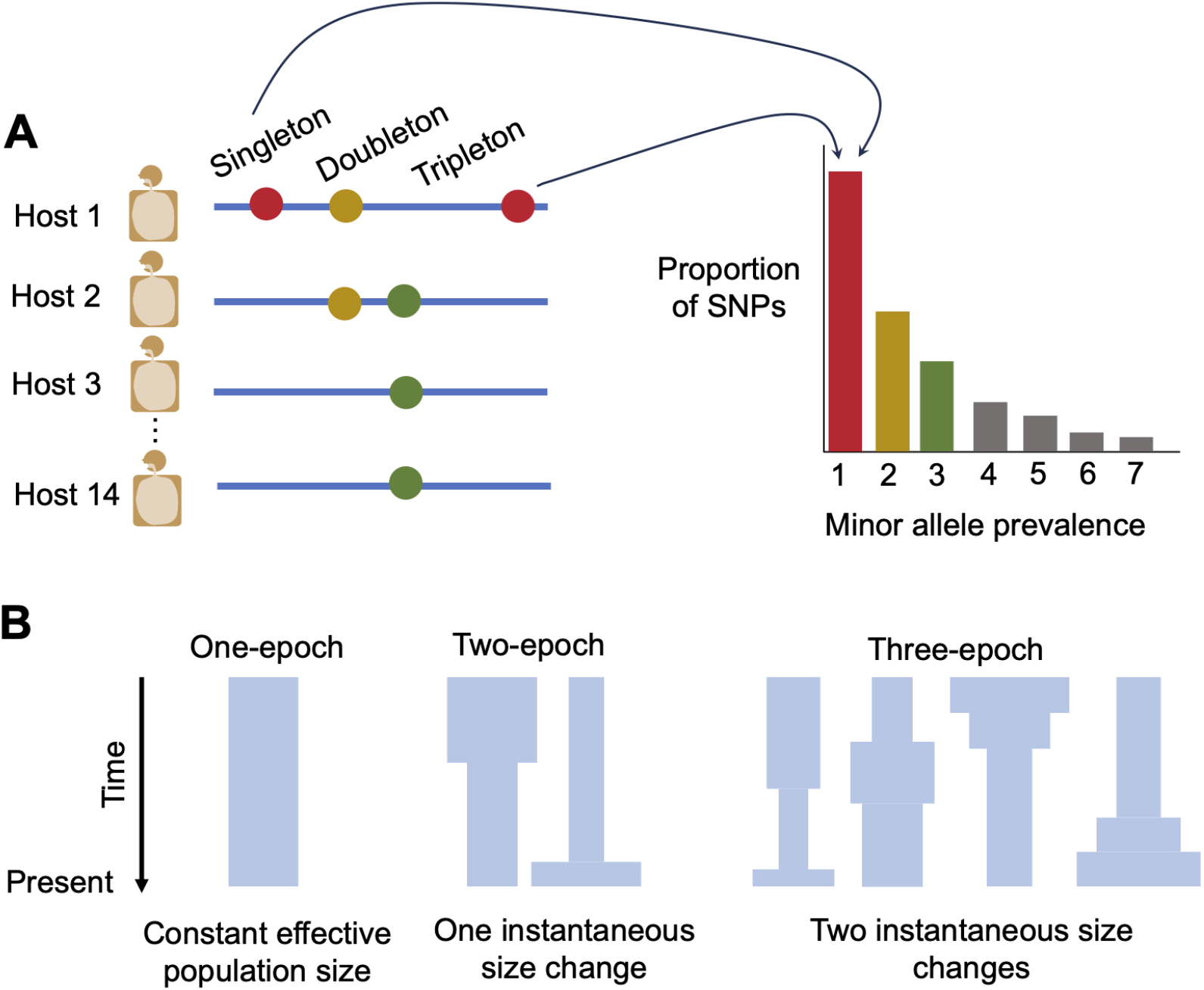
Data and schematic of putative demographic models. (**A**) A single quasi-phased haplotype was inferred for each species for each host with that species. Shown are four example quasi-phased haplotypes. These haplotypes harbor example SNPs that are either ‘singletons’ (present in a single host), ‘doubletons’, ‘tripletons’, etc. SFSs were then constructed by binning the prevalences of alleles found across the quasi-phaseable haplotypes for each species. (**B**) SFSs were then used to infer parameters for one, two, and three epoch models, as depicted. Time is shown on the y-axis. The width of each model represents effective population size at the corresponding time. A one-epoch model has a constant effective population size and can be considered to be a null model against any models which permit effective population size changes. A two-epoch model has a single instantaneous change in effective population size, allowing for inference of contractions or expansions. A three-epoch model has two instantaneous changes in effective population size, allowing for inference of multiple contractions and/or expansions.

To obtain a sample of lineages across hosts, we leveraged a previous approach we developed to ‘quasi-phase’ haplotypes corresponding likely to the dominant strain present in each host for each species (Garud and Good et al. 2019, PLoS Biology; Methods). SFSs were then composed of 1 quasi-phased genome per host for each of 27 of the most prevalent bacterial species present in at least 14 North American gut microbiomes (Methods) (**Figure 2A**).

Since population structure can result in false inferences by generating an excess of SNPs at intermediate frequencies arising from SNPs unique to each subpopulation (Gazave et al. 2014, PNAS; Stadler et al. 2009, Genetic), we controlled for population structure at two levels. First, while different hosts typically harbor their own, genetically distinct set of strains, occasionally there are some exceptionally closely related lineages (divergence < 2 x 10^-4^) circulating in the population (Garud and Good et al. 2019, PLoS Biology). To avoid structure arising from potential clonal expansions, we excluded lineages with divergences < 2 x 10^-4^ to any other lineage in the dataset, as we previously did to control for population structure (Garud and Good et al. 2019, PLoS Biology) (Methods).

Second, previous work has found that several bacterial species present across North American gut microbiomes comprise deeply diverged clades lacking gene flow between groups (Costea et al. 2017, Molecular Systems Biology; Garud and Good et al. 2019, PLoS Biology). Confirming this, lineages belonging to the same clade exhibit significantly higher amounts of recombination than lineages part of different clades, as demonstrated by more rapid rates of LD decay and increased phylogenetic inconsistencies of individual SNPs with a genome-wide phylogeny (Garud and Good et al. 2019, PLoS Biology; Liu and Good 2023, BiorXiv). Thus, to control for population structure as a potential confounder for demographic inference, we analyzed SFSs belonging to lineages comprising the largest clade for each species, based on our previous clade inferences (Garud and Good et al. 2019, PLoS Biology). In **Figure S2** we show the impact of population structure control and subsequent projection on the shape of the empirical SFS.

Finally, selective forces acting on nonsynonymous sites may confound inference of demography by changing the shape or skew of the SFS. For example, purifying selection typically results in an excess of rare variants, thus skewing the nonsynonymous SFS towards low frequency SNPs. By contrast, the SFS of synonymous variants is thought to be more neutrally evolving, thereby allowing for more sensitive inference of the underlying demographic history (Williamson et al 2005, PNAS; Boyko et al 2008, PLoS Genetics). To control for the potential effects of selection, we constructed SFSs using only synonymous SNPs. We initially focused our analysis on the core genome of each species, i.e., genes present in at least 95% of samples (Methods).

#### Demographic inferences with ∂α∂*i*

To infer the demographic histories, we used a statistical inference package known as ∂α∂*i* (Gutenkunst et al. 2009, PLoS Genetics), which is a maximum likelihood inference method based on the SFS. We explored three different putative demographic models in this framework for each species, permitting the inference of population expansions versus contractions **(Figure 2B)**. Selection of the best-fit model was performed using the Akaike information criterion (AIC) (Methods; **Figure S1**).

Demographic inference performed via ∂α∂*i* recovers two main evolutionary parameters: ν, which is the ratio of current effective population size (N_Curr_) to ancestral effective population size (N_Anc_), and τ, the time since the most recent instantaneous size change in units of generations scaled by 2N_Anc_ (Gutenkunst et al. 2009, PLoS Genetics). For each species, we found the maximum likelihood estimates of ν and τ by computing the expected SFS under a given set of parameters and evaluating its fit to the observed SFS (Methods, **Figure 3C-D, Figure S3**).

**Figure 3.**
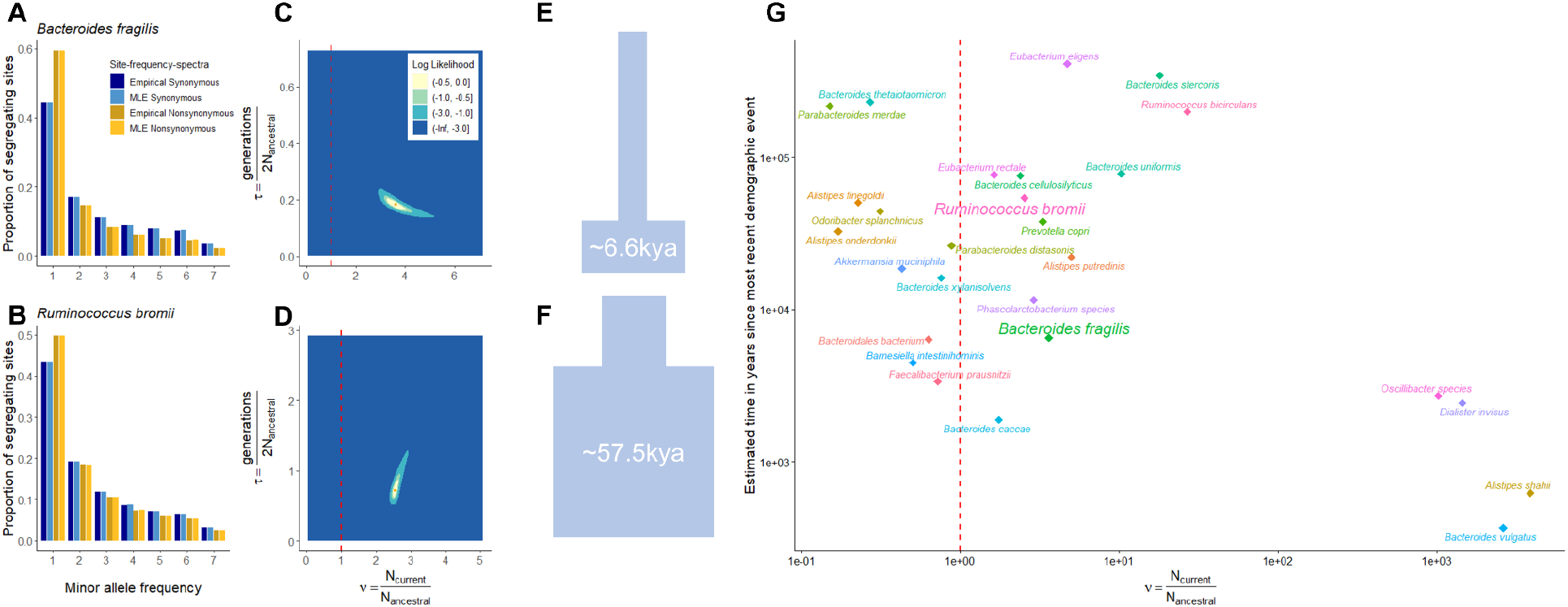
Demographic inference of common commensal gut microbes. Site frequency spectra from the empirical data are compared to those predicted from the maximum likelihood demographic models for two species: (**A**) *Bacteroides fragilis* and (**B**) *Ruminococcus bromii.* “MLE synonymous” shows the expected SFS produced by the ML demographic parameter estimates. “MLE nonsynonymous” shows the expected SFS produced by the ML demographic and selection parameters from a gamma-distributed DFE. 2-dimensional likelihood surfaces of population size, ν (in units of N_Anc_) and time since the onset of the most recent demographic event, τ (in units of 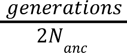) are shown for (**C**) *B. fragilis*, and **(D)** *R. bromii*. A dashed red line separates the parameter space between contractions and expansions. The maximum-likelihood demographic parameters are shown with an orange dot, and colored intervals denote the decrease in log-likelihood from the MLE. The light cyan regions (LL - 3) correspond to the asymptotic 95% confidence interval from the log-likelihood surface. See **Figure S3** for similar plots for all species analyzed. Cartoon schematics (approximately to scale) describing the inferred demography of (**E**) *B. fragilis*, and (**F**) *R. bromii*. **(G)** A summarizing 2-dimensional scatterplot of ν and approximate time in years for ML demographic parameters for each species analyzed in this paper.

As an example, we highlight the demographic inference of two species, *Bacteroides fragilis* **(Figure 3A)** and *Ruminococcus bromii* (**Figure 3B**). For both species, a two-epoch demographic model best fits the data. Maximum likelihood analysis of the SFS of *Bacteroides fragilis* reveals an effective population size expansion in the range of 2.92 to 5.17 times the ancestral effective population size on a time-scale of roughly ∼5,000 to ∼8,000 years ago (these represent the respective lower and upper bounds of ν and τ rescaled to years from the 95% confidence interval from the likelihood surface, see **Figure S4**; Methods). By contrast, the maximum likelihood analysis of the SFS of *R. bromii* is consistent with a population expansion between 2.39 to 2.92 times the ancestral effective population size on a time-scale of roughly ∼40,000 – 100,000 years ago. Confirming these inferences, qualitatively, the observed SFS of both *Bacteroides fragilis* and *R. bromii* have an excess of rare variants compared to a one-epoch model, consistent with an expansion in effective population size.

Across the 27 species for which we were able to infer demographic models, we found that the two-epoch demographic model best fits the data in all analyzed bacteria except for *Odoribacter splanchnicus*, which was best fit to a one-epoch model (**Table S3**). Across species, ν ranges from ∼0.15 to ∼3855, indicating that some bacterial species have experienced a contraction (n=11) while others have experienced an expansion (n=16). In the four cases where ν is extremely large (>100 for the species *Oscillibacter species, Dialister invisus, Alistipes shahii, and Bacteroides vulgatus*), the likelihood surface is very flat and confidence intervals exceptionally large, indicating substantial uncertainty in such large estimate of ν for these species (**Figure S4** and **Table S4**). Finally, when converted to years (Methods), estimates for the timing of population size changes approximately fall between 3.7 x 10^2^ to 4.1 x 10^5^ years ago (**Table S4**). This range of timescales span many important periods of human history, including industrialization (10^2^ years ago), the transition to agriculture (10^4^ years ago), and out of Africa migrations (10^4^-10^5^ years ago).

Together, these findings suggest that commensal gut microbial populations have complex and diverse demographic histories, with some species experiencing substantial population size changes and others exhibiting more stable trends. Additionally, the approximate historical timescales of population demographic changes have occurred over periods that overlap with key events in human history, raising several interesting questions about how human behaviors may have selected for or against certain gut commensal microbiota (Discussion).

### DFEs of gut commensal microbiota display phylogenetic trends

Given inferences of demographic history, we next turned our attention to inferring selection. As seen in **Figure 3A-B**, the observed SFS of synonymous SNPs differs substantially from that of nonsynonymous SNPs. Concretely, **Figure 3A-B** reveals a higher proportion of singletons found among nonsynonymous SNPs compared to putatively neutral synonymous SNPs. This suggests the presence of strong purifying selection acting genome-wide (Bustamante et al. 2001, Genetics; Williamson et al. 2005, PNAS; Eyre-Walker et al. 2006, Genetics; Eyre-Walker and Keightley 2007, Nature Reviews Genetics; Boyko et al. 2008, PLoS Genetics), consistent with other bacterial populations (Hughes 2005, Genetics; Cornejo et al. 2013, MBE). Towards this end, we inferred the distribution of fitness effects (DFE) of nonsynonymous mutations in the 27 common commensal bacterial species for which we inferred demographic models (**Figure 4**).

**Figure 4:**
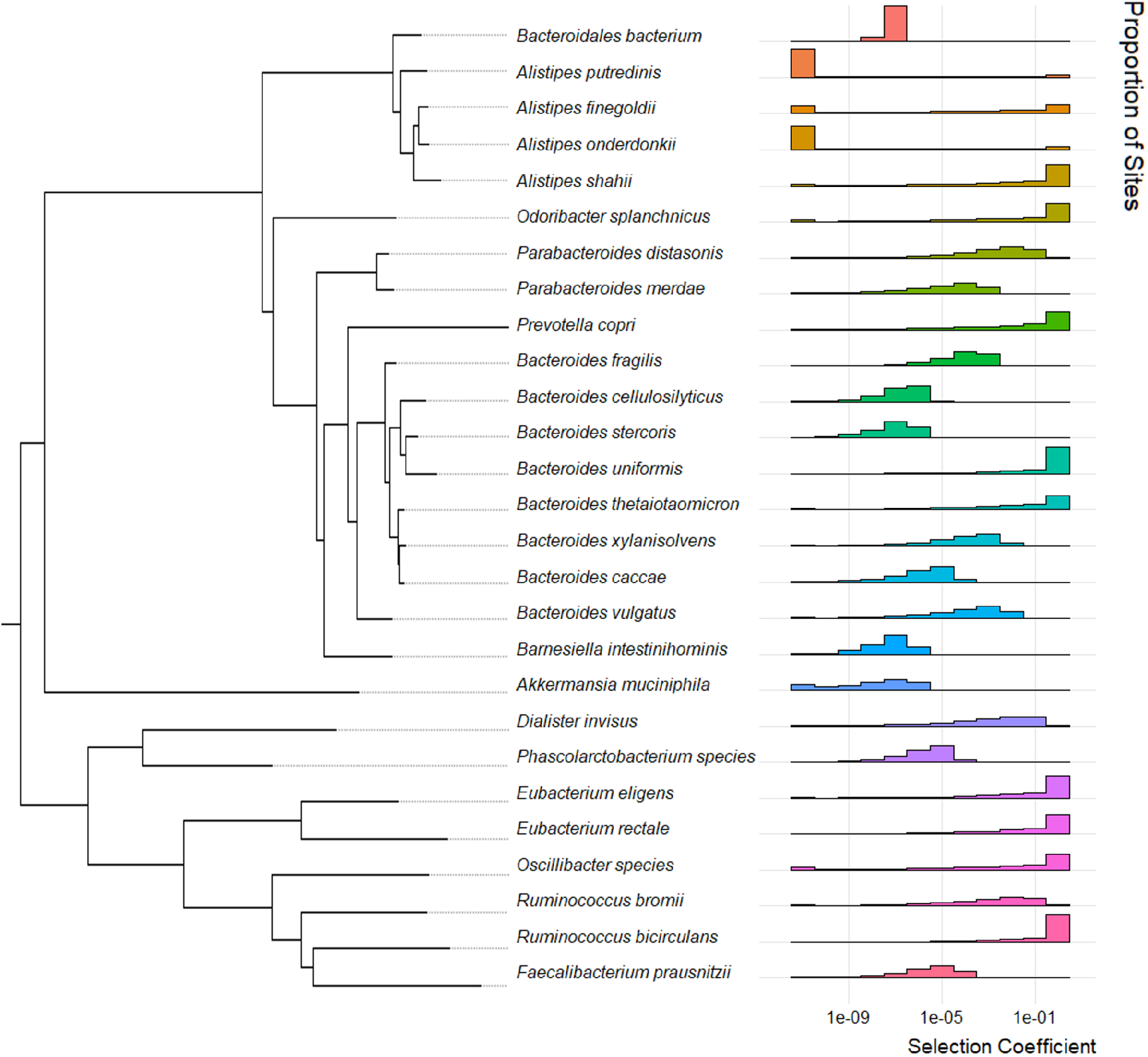
Phylogenetically sorted distributions of fitness effects. Phylogenetically sorted panel of inferred maximum likelihood distributions of fitness effects per species under a gamma-distribution. For each species, the x-axis denotes the log_10_ scaled discrete bin of selective effect, while the y-axis denotes the proportion of sites which fall into each bin.

To infer the DFE, we constructed SFSs for nonsynonymous SNPs from the same species for which we constructed SFSs composed of synonymous SNPs. We then used Fit∂α∂*i* (Kim et al. 2017, Genetics) to infer the DFE of nonsynonymous mutations based on this SFS. As demographic history can affect the SFS, Fit∂α∂*i* infers selection conditional on the demographic history that we inferred from the synonymous SFS.

We examined the fit of several functional forms of the DFE to the SFS (Kim et al. 2017, Genetics). One such functional form is a gamma distribution, characterized by flexible parameters that generate diverse distributions. Another such functional form is the neu+gamma distribution, which is a mixture distribution of a gamma distribution and a point mass on neutrality, i.e., a proportion of mutations which are neutral. The neu+gamma distribution has one additional parameter compared to the gamma distribution, comprising the proportion of neutral or non-deleterious mutations. The model parameters that produce a nonsynonymous SFS most closely resembling the observed nonsynonymous SFS are considered representative of the underlying evolutionary parameters. We compared the AIC of the gamma-distributed DFE (**Figure 4**) with the neu+gamma distributed DFE (**Figure S5**), and found that there was no significant improvement in fit for the neu+gamma compared to the gamma, except in the case of a single species, *Prevotella copri*. Given a general absence of improved model fit for the neu+gamma model over the gamma model, below we further analyze the gamma-distributed DFEs (**Table S5**).

Fitting a gamma model to the SFS of nonsynonymous variants provided insights into the varying degree of deleterious selection coefficients (*s*) expected for each species. For example, in *R. bromii*, we inferred that 16.2% of non-synonymous mutations were weakly deleterious or neutral (*s* < 10^-6^), 49.2% were moderately deleterious (10^-6^ < *s* < 10^-2^), 33.7% were highly deleterious (10^-2^ < *s* < 0.5), and 0.1% were lethal (*s* ≥ 0.5). By contrast, in *B. fragilis*, we inferred that 11.3% of mutations were weakly deleterious or neutral (*s* < 10^-6^), and 88.7% were moderately deleterious (10^-6^ < *s* < 10^-2^), while no mutations in *B. fragilis* were found to be highly deleterious (10^-2^ < *s* < 0.5) or lethal (*s* ≥ 0.5).

We phylogenetically ordered (Methods) the 27 inferred gamma-distributed DFEs (**Figure 4**) and neu+gamma-distributed DFEs (**Figure S5**). This visualization reveals some evidence of phylogenetic trends in the DFE of the gut microbiome. For example, both species in the *Eubacterium* genus were inferred to have DFEs skewed towards highly deleterious and lethal mutations (on average, approximately 75% of new mutations are inferred to have *s* > 10^-2^). By contrast, the majority of species in the *Bacteroides* genus have DFEs skewed towards weakly or moderately deleterious mutations (on average, about 82% of sites are inferred with *s* < 10^-2^). While broad trends across well-represented genera are present, there also exist departures. For example, although the vast majority of new nonsynonymous mutations in the *Bacteroides* genus are weakly or moderately deleterious, about 65% of mutations in *B. uniformis* and 32% of new mutations in *B. thetaiotaomicron* are lethal, with *s ≥* 0.5.

To quantify whether gamma-distributed DFEs between all pairs of the 27 species in our dataset are distinct, we implemented a likelihood ratio test similar to that in Huber et al. 2017 (Methods). Specifically, we compared the log-likelihood of two models: (1) a full model where pairs of species have independently inferred shape and scale parameters and (2) a constrained model in which pairs of species have the same shape and scale parameters.

We performed the likelihood ratio test using two different null models: a null model in which we expect that the distribution of selection coefficients, *s*, is the same between pairs of species (**Figure S6A**), and a null model in which we assume the distribution of population-scaled selection coefficients, 2N_Anc_*s*, is the same between pairs of species (**Figure S6B**). Comparing the selection coefficient, *s,* across species gives insight into the relative fitness of individual mutations, whereas comparing 2N_Anc_*s* across species gives insight into how selection and drift act together.

The likelihood ratio test, when evaluating with *s*, indicates that the DFE is significantly different in 131 out of 351 (about 37%) pairs of species, after Bonferroni correction. When evaluating with 2N_Anc_*s,* the likelihood ratio test is significant in 206 pairs (about 59%), after Bonferroni correction (**Figure S6**). These results suggest that bacterial DFEs differ quite substantially across species. By contrast, comparisons within genera reveal some evidence of phylogenetic congruence of the DFE. Concretely, of 131 significantly different DFEs using a null model where *s* is assumed equivalent between pairs of species, 118 (about 90%) of such tests were between species of differing genera. In the same vein, of 206 significantly different DFEs using a null model where 2N_Anc_*s*, is assumed equivalent between pairs of species, 191 (about 93%) of such tests were between species of different genera. These results suggest that phylogenetically related species have more similar DFEs.

### Accessory genes experience more drift compared to core genes

Thus far, our analyses only have considered core genes present in ≥95% of samples. Next, we evaluated whether demography and selection may have acted differently on accessory genes present in 30-70% of samples as opposed to core genes.

By visual inspection, the synonymous SFSs constructed from the accessory genes exhibit a proportional depletion of rare variants relative to the core genome of the same species (**Figure S7**), consistent with possible contractions in effective population size due to demography and/or linked selection. To more specifically investigate if accessory versus core genes display distinct demographic histories or if selection acts differently, we first fit demographic models and distributions of fitness effects for accessory gene SFSs. Due to data paucity (e.g. insufficient numbers of base pairs represented in the SFS), we were only able to find well-fitting models for 7 out of 27 species (**Figure 5C, Figure S8**). For these 7 species, we found that two-epoch demographic models best fit the data (**Table S6**), and that on average ν and N_Anc_ are smaller for accessory genes relative to core genes (**Table S6**). We recapitulate this result in 23 species when inferring the demography from SFSs constructed from both core and accessory genes and find that there is a systematic decrease in ν relative to when demographic inferences are performed only from core genes (P-value=8.26 x 10^-3^, Wilcoxon rank sum test, **Figure S9**).

**Figure 5:**
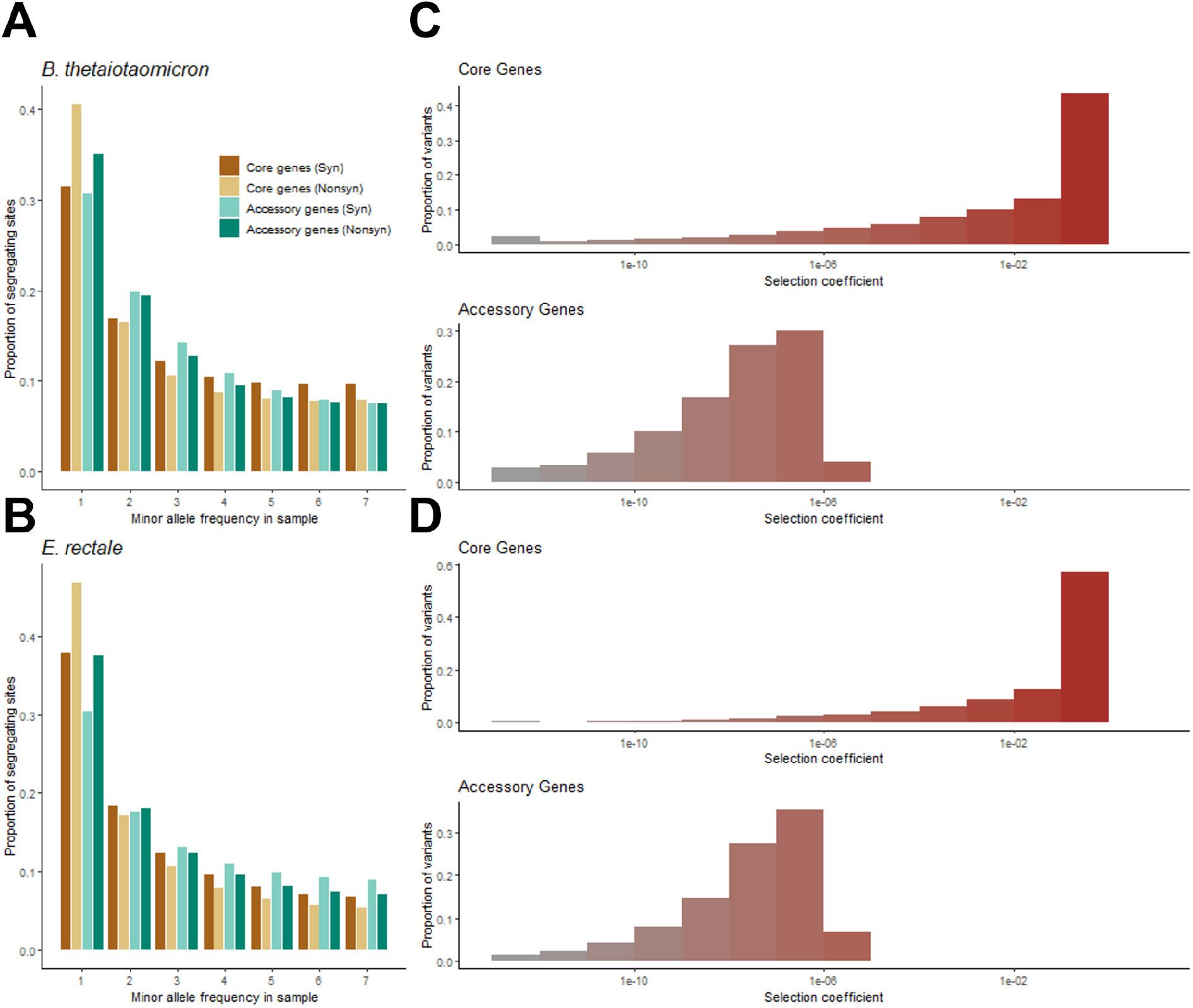
Differences in selective effects between core and accessory genes. Empirical site frequency spectra of synonymous and nonsynonymous SNPs from core genes and accessory genes for two species: (**A**) *B. thetaiotaomicron* and (**B**) *Eubacterium rectale*. The maximum likelihood inferred gamma-distributed DFEs of *s* are shown for (**C**) *B. thetaiotaomicron* and (**D**) *E. rectale*. For each species, the x-axis denotes the log10 scaled discrete bin of selective effect, while the y-axis denotes the proportion of sites which fall into each bin. Bins are color coded from gray to dark red according to the inferred lethality on the x-axis. See **Figure S10** for similar plots for all species analyzed.

These results are consistent with accessory genes being present in fewer host microbiomes. We note that we excluded any gene crossing species boundaries or has the potential to, so as to avoid spurious read mapping (**Methods**). However, had we included these genes, Ne values for accessory genes might have been larger given their broader prevalence. Therefore it is possible that by restricting our focus to genes present only within a species, the lower Nanc could be due to gene loss in the accessory genome.

Next we inferred the DFE for accessory genes for the 7 species with well-fitting demographic models (**Table S7**). Confirming our visual comparison of the nonsynonymous SFS from core vs. accessory genes (**Figure 5A-B**) in which there is again a depletion of rare variants for accessory genes, the DFEs of accessory genes are less deleterious than the DFE of core genes (**Figure 5C-D**). We performed a likelihood ratio test (Methods) of similarity between the DFE of accessory genes vs. the DFE of core genes within each species (**Table S8**). When using a null model in which *s* is assumed constant between the accessory and core genes, we reject the null hypothesis for only 1 species, *B. vulgatus*. However, when using a null model in which 2N_Anc_*s* is assumed equivalent between the accessory and core genes, we reject the null hypothesis for 5 out of 7 species, after Bonferroni correction. Taken together, these results suggest that differences in effective population size between core and accessory genes largely drive differences in their SFSs, consistent with the smaller N_Anc_ for accessory genes inferred above. However, as there is limited power to detect differences in *s* between core and accessory genes due to paucity of SNPs in the accessory genes, we cannot fully rule out subtle shifts in the selection effects of individual mutations.

## DISCUSSION

In recent years, much has been learned about the evolutionary dynamics of human gut microbiota, both within and across hosts. Nevertheless, there remains a need to comprehensively investigate how demographic history and selective forces influence genetic variation across commensal gut populations (Costea et al. 2017, Molecular systems biology; Truong et al. 2017, Genome Research; Garud and Pollard 2020, Trends in Genetics; Dapa et al. 2023, Current Opinion). This study represents the first systematic examination of the demographic histories and distribution of fitness effects (DFEs) of commensal gut bacteria across hosts. To infer demographic histories and DFEs for 27 species, we used a site frequency spectrum-based analysis, an approach commonly used in eukaryotes (Beichman et al 2018, Annual Reviews; Gutenkunst et al 2009, PLoS Genetics) but has been under-utilized for bacterial populations (Cornejo et al. 2013, MBE; Pepperell et al 2013, PLoS Pathogens). We found that commensal microbiota have experienced a range of population size changes over the course of human history, and we inferred finer distributions of fitness effects compared to previous work, which estimated the proportion of sites experiencing a single selection coefficient rather than a range of coefficients. (Garud and Good et al. 2019, PLoS Biology; Cornejo et al. 2013, MBE). Finally, we found that closely related species exhibit more similar DFEs than those from different clades, and that accessory genes have smaller effective population sizes, reducing the potential impact of natural selection.

In our inferences, we assume that the SFS for synonymous SNPs reflects demographic history. However, a myriad of evolutionary processes can affect the SFS (Braverman et al. 1995, Genetics; Lapierre et al. 2017, Genetics). For example, purifying selection can reduce diversity and skew the SFS towards rare variants (Braverman 1995, Genetics; Lapierre et al. 2017, Genetics; O’Fallon et al. 2010, MBE) and selective sweeps can also result in an excess of rare variants (Fay and Wu 2000, Genetics; Braverman 1995, Genetics), both of which would result in false inferences of expansions. While the pervasiveness of purifying selection among commensal bacteria has been well-documented (Schloissnig et al. 2013, Nature; He et al. 2010, PNAS; Garud and Good et al. 2019, PLoS Biology; Zhao, Lieberman et al. 2019, Cell Host Microbe; Rocha and Feil 2010, PLoS Genetics), the extent of selective sweeps across host microbiomes is yet to be quantified. Additionally, while recombination in human gut microbiomes seems to be common and frequent, understanding finer-scale variation of recombination rates along microbial genomes and its impact on linked effects between non-neutral and neutral sites will be important.

Similarly, false population contractions may be inferred if there is some force that inflates intermediate frequency synonymous variants, such as balancing selection or cryptic population substructure (Voight and Pritchard 2005, PLoS Genetics; LaPierre et al. 2017, Genetics). Balancing selection has been inferred using Tajima’s D from a subset of genes but not genome-wide (Moeller 2021, Genome Biology and Evolution). Thus, it is highly unlikely that the signals observed here for population contractions stem from balancing selection as we use genome-wide polymorphism data. While we controlled for population substructure (**Figure S2**, Methods), there still could be residual cryptic population structure that is confounding analyses. Future work investigating the impact of downsampling on the inference of complex demographic models will be an informative avenue of future research.

Despite the simplicity of our models, basic aspects of our inferred demographic models concur with previous estimates of Ne made in the literature. We found that MLEs for N_Curr_ typically fall between 10^6^ – 10^8^, in agreement with a previous study that estimated N_Curr_ using dN/dS to be in the range of 10^6^ – 10^9^ for >150 commensal, pathogenic, and free-living bacterial species (Bobay and Ochman 2018), indicating a general agreement of our results and previous inferences made with different approaches. Four of the species that we analyzed have N_Curr_ on the order of 10^10^, but the likelihood surfaces for these species are extremely flat, yielding almost equivalent log likelihoods for a wide range of values as low as 10^8^ (**Table S4, Figure S4**).

It is striking that many of the demographic changes we infer coincide with key events in human history, such as the industrial revolution, the transition to agriculture, and human migration out of Africa. Even with a lower assumed mutation rate of 10^-10^ per base pair per generation (Barrick and Lenski 2013, Nature Reviews Genetics) (**Methods**), our estimate of time in years still would span these significant human epochs. It is tempting to speculate that events like the agricultural revolution may have been causal of changes in effective population sizes among gut commensals. For example, Cornejo et al. (2013) inferred a population expansion for the cavity-causing pathogen *Streptococcus mutans* ∼10,000 to 20,000 years ago, potentially due to new available ecological niches opening up as a consequence of changes in diet during the agricultural revolution. Indeed, the direction of inferred population size changes concur with changes in relative abundances for some species in industrialized vs traditional populations (e.g. we infer expansions for 6 of 8 *Bacteroides* species, that have been previously shown to be higher in abundance in industrialized populations due to dietary shifts (Sonnenberg and Sonnenberg 2019, Science)).

However, there are some cases where the inferred population size changes do not align with expectations arising from prevalence and abundances in industrialized versus traditional populations. For example we inferred a population expansion for *Prevotella copri* despite it being at markedly lower prevalence and abundance in industrialized versus traditional populations due to reductions in plant-based carbohydrates in western diets (Sonnenberg et al. 2016, Nature; Wu et al. 2011, Science, Tett et al. 2019, Cell Host Microbe, Sonnenberg and Sonnenberg 2019, Science). However, a three-epoch demographic model for *P. copri* has an AIC only ∼5 units higher (**Figure S1**) than a two-epoch model, and in fact does display a severe bottleneck approximately 20,000 – 40,000 years ago followed by a moderate expansion in effective population size in the time since. Such a model is simultaneously consistent with overall reductions in nucleotide diversity observed in North Americans compared to rural Africans (**Figure 1**) and the expansion inferred with the two-epoch model. In general, we find that overall the three-epoch models for most species have very similar AICs and LLs as the two-epoch models, and so it is difficult to distinguish between these models (**Figure S1**). One-epoch models (models lacking any population size change), however, have significantly worse AICs and thus are easier to reject. Future research will be needed to infer more sophisticated models for each of the species in our dataset, as our models are simple and likely do not reflect the full set of demographic changes that have transpired for each species.

Beyond inferring a range of demographic histories across species, we also found differences in their DFEs. Interestingly, we found that phylogenetically related species have more similar DFEs than distantly related species (**Figure S6**) potentially implying functional similarities among closely related species. We note that both differences in *s* and N_Anc_ could be driving differences in the DFE across species, though we find more differences between pairs of species when comparing the DFE in terms of *s* rather than 2N_Anc_*s*, reflecting potential shifts in the deleteriousness of individual mutations in different species. By contrast, the differences in the DFEs for core vs accessory genes are less likely due to differences in *s* and instead due to differences in N_Anc_. In support of this, accessory genes have lower N_Anc_ than core genes (**Figure S13**), and have elevated levels of pN/pS (N’Guessan et al. 2021, Genome Biology and Evolution; Cooper et al. 2010, PLoS Computational Biology; Olm et al. 2021, Nature Biotechnology), suggesting that weakly deleterious mutations can drift to higher frequency due to the smaller effective population size.

Finally, it is worth noting that our DFE and demographic history inferences rely on genotypes supported by a majority of reads within hosts (Methods), mitigating the impact of sequencing errors on our inferences. Consequently, our inferred DFEs represent what could be considered an ’average’ across hosts and may not fully capture within-host DFE variations, which can potentially differ among individuals and environments (Dapa et al. 2023, Current Opinion). Future research utilizing experimental evolution techniques (Shiver 2023, BioRxiv; Wong and Good 2022, BioRxiv) will be pivotal in unraveling differences between within and across-host DFEs and their associations with dynamic processes within human microbiomes and FMT outcomes.

In summary, this work advances our understanding of the evolutionary dynamics of human gut microbiota, highlighting the influence of demographic history and selective forces on genetic variation. Our findings underscore the complexity of these interactions and their implications for the human microbiome, which future work will undoubtedly further elucidate.

## ACKNOWLEDGEMENTS

This work was funded in part through the NIGMS NIH T32 5T32GM008185 (Systems in Integrative Biology) training grant to JM and NIGMS NIH awards R35GM151023 to NRG and R35GM119856 to KEL. NRG also received support from an NSF CAREER award (no. 2240098), a Paul Allen Research Foundation grant, the UC Hellman Fellows grant and a UCLA faculty research grant. The authors would like to thank Daisy Chen and Richard Wolff for their computational help, Jesse Shapiro and Pleuni Pennings for reading the manuscript and providing feedback, and Benjamin Good, Emily Ebel, and all members of the Garud and Lohmueller labs for helpful discussions.

## CODE AVAILABILITY

All necessary metadata, as well as the source code for the bioinformatics pipeline, downstream analysis, and figure generation are available at Github: https://github.com/garudlab/microbiome_demography_manuscript

## METHODS

### Data

Raw whole-genome shotgun sequencing reads for North American metagenomic samples analyzed in this study were downloaded from the Human Microbiome Project Consortium (2012); Lloyd-Price et al. (2017) (URL: https://aws.amazon.com/datasets/human-microbiome-project/). African metagenomic samples from Pasolli et al. (2019) were downloaded from ebi.ac.uk with accession number PRJNA485056. Accession numbers associated with this data are in **Table S1** and **Table S2**.

For North American gut microbiomes, there were 471 fecal samples available from 250 healthy North American individuals. A subset of these 250 individuals were sampled at 2 or 3 time points. We analyzed data from a single time point per host per species in the North American dataset. Additionally, since previous work has shown that technical and sample replicates of the same fecal sample have little genomic variability (Lloyd Price et al. 2017, Nature; Nayfach et al. 2016, Genome Research; Maghini et al. 2023, Nature Biotechnology), we merged FASTQ files for technical and sample replicates from the same time point to increase coverage as in Garud Good et al. 2019, PLoS Biology. For Madagascar gut microbiomes, we analyzed all 112 fecal samples from 112 healthy Madagascar individuals (**Table S1**, **Table S2**).

### Identification of bacterial species, genes, and SNVs

We used a standard reference-based approach called MIDAS to quantify bacterial species abundances and gene and SNV content (Nayfach et al. 2016, Genome Research).

The first step of the MIDAS pipeline is to determine the species present in each set of sample(s) for each host. MIDAS quantifies the relative abundance of species in a given sample by mapping sequencing reads to a reference database of single-copy “marker” genes unique to each species. We used database version 1.2, downloaded on November 21^st^, 2016 (Nayfach et al. 2016, Genome Research). A species was considered present if it had an average marker gene coverage of at least 3x for a given sample.

Second, MIDAS quantifies SNVs for each species for each sample. Similar to the “species” step, this “SNV” step leverages a standard reference-based approach in which reads are mapped to a single genome per species. To avoid reads mapping spuriously to species not present in the dataset, reads were mapped only to species truly present in the sample as per the species step defined above. Mapping was done with Bowtie2 (Langmead and Salzberg 2012, Nature Methods) with the following default MIDAS mapping thresholds: global alignment, MAPID ≥ 94.0%, READQ ≥ 20, ALN_COV ≥ 0.75, and MAP ≥ 20.

MIDAS annotates SNVs as either 1D or 4D, indicating codon degeneracy. For example, 1D means that any nucleotide change will result in an amino acid change, resulting in a nonsynonymous site. By contrast, 4D means that any nucleotide change will result in a synonymous site.

Finally, MIDAS identifies gene copy number variants by aligning reads to a pangenome, i.e., the union of all genes found in any strain present in the MIDAS database for a given species. Once again, reads were mapped to pangenomes only for those species considered to be present. Standard MIDAS read coverage parameters were used for calling copy number variants and as described in Garud and Good et al. 2019 (PLoS Biology).

Since there may be orthologs of the same gene present in multiple species, these genes can potentially result in spurious read donating and recruiting. Therefore, any gene with ≥ 95% ANI between different species’ pangenomes was excluded from further analysis. Genes with ≥ 95% ANI with another gene in multiple species’ pangenomes were previously identified in Garud and Good et al. 2019 (PLoS Biology).

To identify which genes to include in our analysis, we considered the copy number of each gene. Genes were included in our data set if their estimated copy number was ≥ 0.3 and ≤ 3.0 (Garud and Good et al. 2019 PLoS Biology). Next, to differentiate between core and accessory genes, we considered the prevalence of each gene: core genes were identified as genes with prevalence ≥ 95%, while accessory genes were identified as genes with prevalence between 30% and 70%.

Once core and accessory genes were identified, sites were included using a minimum sample depth of 20X coverage, and a minimum site depth of 20 reads.

### Estimation of nucleotide diversity

Within-host nucleotide diversity (π*_within_*) of each species was estimated as:

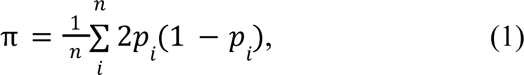

where *p*_*i*_ is the frequency of the reference allele at site *i* and *n* is the length in base pairs of the core genome (Gillespie (2004).

Between-host nucleotide diversity (π*_between_*) was computed similarly. The allele frequency at a given site *i* in the pooled population was first computed as:

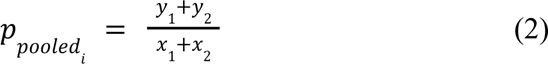

where *y*_1_ and *y*_2_ correspond to the number of reads supporting the reference allele in samples 1 and 2, respectively, and *x*_1_ and *x*_2_ correspond to the total number of reads at the given site in samples 1 and 2, respectively. Π_*between*_ was then computed as for each pair of metagenomic samples with equation (1) using *p* instead of *p*_*pooledi*_. All values of π_*between*_ across pairs of hosts were then averaged for each species, yielding a species-specific estimate of between-host nucleotide diversity.

### Quasi-phasing

To estimate quasi-phased haplotypes from individual hosts, we identified hosts in which the lineage structure of their bacteria was simple enough to assign alleles to the dominant lineage with high confidence as described in Garud and Good et al. 2019 (PLoS Biology). Since both our paper and Garud and Good et al. 2019 (PLoS Biology) utilize the same HMP data for analyses, we used the same sample x species identified in Garud and Good et al. 2019 (PLoS Biology) that were classified as quasi-phaseable. We then estimated the genotype of the dominant lineage by assigning the major allele of each site only if it had a frequency of 0.8 or greater. Any site with a major allele frequency between 0.5 and 0.8 was considered missing data. The number of quasi-phaseable samples for each species present in the largest clade is shown in **Figure S12**.

### Phylogenetic ordering

To order species phylogenetically (such as in **Figure 4**), we used the phylogenetic tree from Nayfach et al. 2016 (Genome Research) and abstracted a subtree of relevant species using the APE package in R (Paradis and Schleip 2019, Bioinformatics).

### Calculating the Site-frequency Spectrum

We compute ‘folded’ site-frequency spectra (SFS) for both synonymous and nonsynonymous sites from quasi-phased bacterial genomes. The folded SFS describes the distribution of minor allele frequencies across the genome. ‘Folded’ indicates that since the ancestral versus derived state of alleles are unknown, any allele frequency (*f*) that is > 50% becomes 1-*f*. Folded SFSs have been shown to yield accurate inferences of the distribution of fitness effects of deleterious mutations (Keightley and Eyre-Walker 2007; Boyko et al. 2008, PLOS Genetics; Kim et al. 2017, Genetics).

The SFS can be represented as a vector, *X* = [*X*_1_, *X*_2_, …, *X*_*n* - 1_] in which each element of the vector describes the number of SNPs at frequency *i* given *n* chromosomes. In other words, *X*_1_ is the number of singletons, *X*_2_ is the number of doubletons, etc. Thus, the SFSs in this paper describe the number of QP samples a SNP appears in.

A site is considered to be variable if at least 1 quasi-phased genome has a different allele from other quasi-phased genomes in our dataset. The population frequencies of alleles can range from 0%, meaning the given site is invariant in the population, to 50%, meaning half the individuals in the sample possess the variant allele. To summarize the distribution of variant allele frequencies, we bin alleles based on their population frequencies.

### Projection of the SFS

The empirical SFS may show irregular peaks and valleys due to missing data (**Figure S2B**), which may confound inference from and modeling of the SFS, as some sites are not accurately called for all individuals. To diminish the confounding effects of missing data, we projected the empirical SFS down from a larger sample size to a smaller sample size of 14 haplotypes for all species. In other words, for all sites present in ≥14 hosts, we computed the expected frequency of each SNP in a sample of 14 haplotypes using a hypergeometric distribution (Gutenkunst et al. 2009, PLoS Genetics). Sites with data in <14 hosts were omitted.

### Inference of demographic models

We consider three model specifications: 1) a one-epoch model consisting of a constant effective population size, 2) a two-epoch model consisting of two epochs, i.e., two periods of time, separated by one effective population size change, and 3) a three-epoch model consisting of three epochs separated by two effective population size changes **(Figure 2**).

We use the statistical inference package, ∂α∂*i* (Gutenkunst et al. 2009, PLoS Genetics) to infer two main demographic parameters for each effective population size change, τ and ν. The time of the population size change is represented by τ, in units of 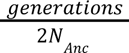, where N_Anc_ is the ancestral effective population size. The parameter ν denotes the ratio of effective population size after the population size change relative to N_Anc_.

To find the maximum likelihood parameters, we defined a grid of points over a likelihood surface parameterized by τ and ν. For a given point on the likelihood surface, *∂a∂i* estimates the model spectrum expected under those evolutionary parameters using diffusion theory. The multinomial log-likelihood function is then used to assess the fit of the model spectrum to the empirical data as described in Gutenkunst et al 2009 (PLoS Genetics). We then performed a gradient-descent based approach to evaluate points along the likelihood surface to find the MLEs. The pair of parameters which yielded the highest log-likelihood were assumed to represent the underlying evolutionary history. We did not force parameter bounds on the likelihood surface, allowing the gradient-descent search to evaluate the entirety of the log likelihood surface. In addition, at least 25 initial parameter guesses, spanning a large range across the parameter surface, were evaluated to ensure that the likelihood surface did not converge at a local maximum specific to a set of starting parameters. We used the chi-square approximation to the log-likelihood ratio to obtain the critical values for the asymptotic 95% confidence intervals of the demographic parameters. The CIs included all parameter values within 3 log-likelihood units (df=2, reflecting the 2 parameters) from the MLE.

### Estimation of ancestral population size

We find N_Anc_ from θ*_s_* as:

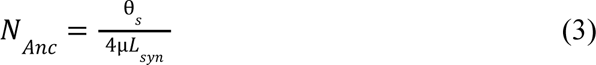

where µ is the mutation rate per site per generation and *L*_*syn*_ is the number of synonymous sites. θ*_s_* is the population-scaled mutation rate, optimized as the scaling factor between the empirical data and the model spectrum which best fits the data. We define µ as having an estimate of 4.08 x 10^-10^ per site per generation, based on experimental estimation for neutral evolution in bacteria (Drake 1991, PNAS).

### Computation of estimated times of demographic size changes

We convert the time parameter, τ, which is in units of 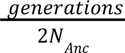, into units of time in years. We chose to assume a conservative generation time of one generation per day, based on estimates of bacterial growth (Savageau 1983, The American Naturalist; Gibbons and Kapsimalis 1967, Journal of Bacteriology; Ghosh and Good 2022, PNAS). Thus, we find:

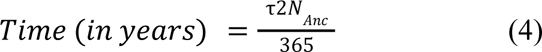

### Inference of the distribution of fitness effects of nonsynonymous mutations

To infer the distribution of fitness effects (DFE), we utilize the site-frequency spectra of nonsynonymous variants. We use a statistical inference package, Fit∂α∂*i*, developed by Kim et al. 2017 (Genetics). Fit∂α∂*i* uses a maximum-likelihood approach to infer the DFE from site-frequency spectra. We note that *s* in Fit∂α∂*i* refers to the change in relative fitness of the heterozygous genotype and is equivalent to the fitness in the haploid.

We examined two different probability distributions for the DFE:a gamma-distributed DFE, and a neutral+gamma-distributed DFE. Under a gamma-distributed DFE, the selective effect of a given mutation, denoted as *s*, is randomly drawn from a gamma distribution, parameterized by *shape*, denoted as α and *scale*, denoted as β. Under a neutral+gamma-distributed DFE, we assume that *s* is drawn from a gamma distribution, but additionally include a parameter for a proportion of sites assumed to be neutral, generating a point-mass at *s=0*, parameterized as *p_neu_*.

The parameters for the DFEs described above were inferred using Fit∂α∂*i*. Importantly, because demography can impact the SFS, we condition on the demographic models described above that we inferred from synonymous variants. Fit∂α∂*i* models the expected nonsynonymous SFS for a given combination of DFE parameters and a demographic history. Then, the fit to the data is computed using a Poisson log-likelihood function. In contrast with the multinomial likelihood, the Poisson likelihood additional requires an *a priori* assumption about the population-scaled mutation rate of nonsynonymous sites, denoted as θ_*ns*_. θ_*ns*_ is obtained by multiplying the the best-fitting θ*_s_* from demographic inference by the expected ratio of nonsynonymous to synonymous mutation rates. We assumed this ratio to be θ_*ns*_/θ_*s*_= 2. 31 (Huber et al. 2017, PNAS). We found the DFE parameters that maximized the log-likelihood by gradient descent over a two-parameter likelihood surface parameterized by α and β. Similar to inference of demography, we did not force parameter bounds on the likelihood surface, allowing the gradient-descent search to evaluate the entirety of the log likelihood surface. Furthermore, at least 25 initial parameter guesses, spanning a large range across the parameter surface, were evaluated to ensure that the likelihood surface did not converge at a local maximum specific to a set of starting parameters.

### Comparing DFEs across species

To quantify the similarities and differences between the DFEs of two species, we performed a likelihood ratio test (LRT) comparing the fit of models of the the DFEs of nonsynonymous mutations assuming that the shape and scale parameters are the same in both species (the null model) species vs. a model where each species can have its own shape and scale parameters (the alternative model).

Consider two species, *A*, and *B*, for which we have independently inferred demographic histories, respectively denoted as Θ_*D*,*A*_ and Θ_*D*,*B*_. Conditional, on these best-fitting demographic histories, we then independently fit the DFE for both species under a gamma-distributed model specification, yielding a shape parameter α and a scale parameter β for each species. To treat species A and B as independent (Boyko et al. 2008, PLOS Genetics, Lawrie and Petrov 2014, Trends in Genetics), we used a Poisson composite likelihood function identical to the likelihood function used in Huber et al. 2017, reproduced here:

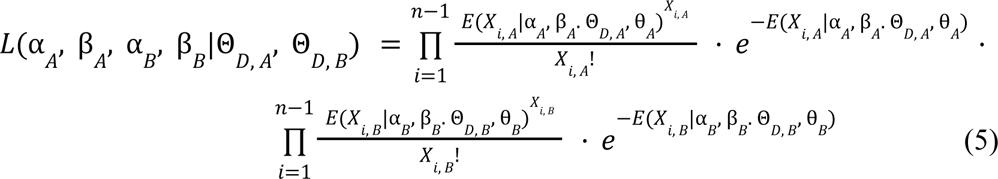

where *n* and *m* denote the sample size of species *A* and *B* respectively, and *X*_*i*, *A*_ denotes the number of SNVs at frequency *i* for a given species, *A*.

To test whether the shape (α) and scale (β) parameters of species *A* differ from those in species B, we utilized the likelihood ratio test similar to that proposed in Huber et al. 2017:

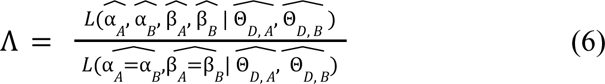

The null hypothesis for this test is the likelihood of the constrained model, i.e., the case in which α_*A*_ = α*_B_* and β_*A*_ = β*_B_*, while the full model allows for α_*A*_ ≠ α*_B_* and β_*A*_ ≠ β*_B_*. We optimized the likelihood function under both the constrained model and the full model. We optimized the constrained model by considering a grid of values for α and β. We then used Fit∂α∂*i* to calculate a site-frequency spectrum of nonsynonymous variants for each pair of parameters for each species, and computed the log likelihood for the fit of each simulation to the data. Then, for each pair of parameter values, we summed the log-likelihood over both species. The pair of parameters with the highest log-likelihood was the MLE for this constrained null model.

Asymptotically, 2Λ follows a χ^2^ distribution with two degrees of freedom, as the full model has two additional free parameters relative to the constrained model. The critical value for the likelihood ratio test is approximately 6 for a critical region of 0.05 for a singular test; however, given that we perform 351 pairwise comparisons, we approximate the critical value as 17.71.

### Comparing DFEs between core and accessory genes

Similar to the method by which we compare DFEs across species, we performed an LRT comparing DFEs of core versus accessory genes, whereby the null hypothesis is a constrained model which assumes that core and accessory genes have the same alpha and beta parameters. The alternative hypothesis is a full model in which alpha and beta may be different for core and accessory genes.

We perform comparisons of the DFE between core and accessory genes for seven species; thus, the critical value for this likelihood ratio test is approximately 9.88, after Bonferroni correction.

## SUPPLEMENT

**Table S1: North American fecal sample accession numbers (see attachment)**

https://github.com/garudlab/microbiome_demography_manuscript/blob/main/Supplement/Supplemental_Table_1.csv

We analyzed 471 fecal metagenomic samples from 250 healthy North American hosts. Listed are the subject identifiers, sample identifiers, run accessions, country of the study performed, continent of the study performed, visit number, and study number (Turnbaugh et al. 2007, Human Microbiome project (2012), Lloyd-Price et al. 2017, Nature).

**Table S2: Madagascar fecal sample accession numbers (see attachment)**

https://github.com/garudlab/microbiome_demography_manuscript/blob/main/Supplement/Suppl emental_Table_2.csv

We analyzed 112 fecal metagenomic samples from 112 healthy African hosts. Listed are the run accessions, sample accessions, experiment accessions, study accessions, and FTP links. The study accession number is PRJNA485056 (Pasolli et al. 2019).

**Table S3: Demographic inference for core genes (see attachment)**

https://github.com/garudlab/microbiome_demography_manuscript/blob/main/Supplement/Supplemental_Table_3.csv

Phylogenetically sorted summarizing table of demographic inference for core genes of 27 common commensal gut microbiota. Three model specifications were considered: a one-epoch demographic model, a two-epoch demographic model, and a three-epoch demographic model (Methods). Listed are the log likelihood, AIC, and maximum likelihood parameters for each species and model specification. τ is converted to time in years as described in the methods. For all species except *Odoribacter splanchnicus*, a two-epoch demographic model best fits the data.

**Table S4: Lower and upper bounds of 95% confidence intervals of ν and time for two-epoch demographic models (see attachment)**

https://github.com/garudlab/microbiome_demography_manuscript/blob/main/Supplement/Supplemental_Table_4.csv

Phylogenetically sorted summarizing table of lower and upper bounds for ν and time in years since the most recent demographic event. Upper and lower bounds were computed using the 95% confidence interval of the log likelihood surface, approximated as parameter space within 3 log likelihoods of the MLE (Figure S1; Methods).

**Table S5: DFE inference for core genes (see attachment)**

https://github.com/garudlab/microbiome_demography_manuscript/blob/main/Supplement/Supplemental_Table_5.csv

Phylogenetically sorted summarizing table of DFE inference for core genes of 27 common commensal gut microbiota. Two model specifications were considered: a gamma-distributed DFE, and a neu+gamma-distributed DFE (Methods). Listed are the log likelihood, AIC, and maximum likelihood parameters for each species and model specification.

**Table S6: Demographic inference for accessory genes (see attachment)**

https://github.com/garudlab/microbiome_demography_manuscript/blob/main/Supplement/Supplemental_Table_6.csv

Phylogenetically sorted summarizing table of demographic inference for accessory genes of 7 common commensal gut microbiota. Listed are the log likelihood, AIC, and maximum likelihood parameters for each species and model specification. τ is converted to time in years as described in the methods. For ease of comparison, the ancestral effective population size inferred from core genes is shown as the right-most three columns.

**Table S7: DFE inference for accessory genes (see attachment)**

https://github.com/garudlab/microbiome_demography_manuscript/blob/main/Supplement/Supplemental_Table_7.csv

Phylogenetically sorted summarizing table of DFE inference for core genes of 7 common commensal gut microbiota. Listed are the log likelihood, AIC, and maximum likelihood parameters for each species and model specification.

**Table S8: LRT statistics for core vs. accessory genes (see attachment)**

https://github.com/garudlab/microbiome_demography_manuscript/blob/main/Supplement/Supplemental_Table_8.csv

Testing the null hypothesis that the DFEs for core and accessory genes from the same species have the same gamma distribution. Two forms of null hypothesis were tested: one in which the selection coefficient *s* is assumed equal between core and accessory genes, and one in which the population-scaled selection coefficient, 2N_Anc_*s* is assumed equal between core and accessory genes (Methods). Given 7 tests, the critical value for statistical significance after Bonferroni correction is approximately 9.88.

---

**Figure S1:**
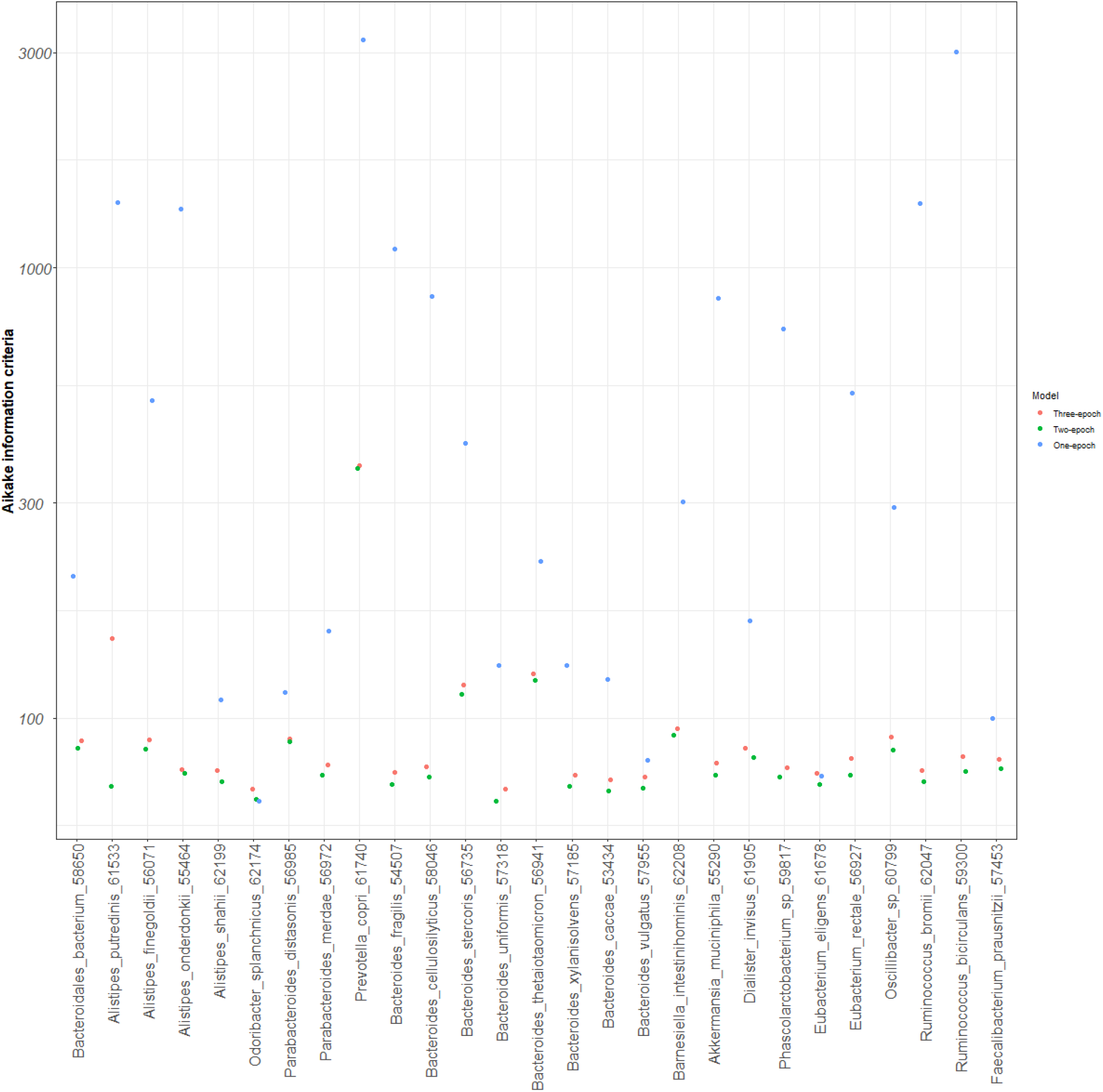
Akaike information criterion for one-epoch, two-epoch, and three-epoch demographic models inferred in this paper. https://github.com/garudlab/microbiome_demography_manuscript/blob/main/Supplement/Supplemental_Figure_1.png Phylogenetically sorted panel of the Akaike information criteria computed for 1, 2, and 3-epoch models inferred in this paper for each species.

**Figure S2:**
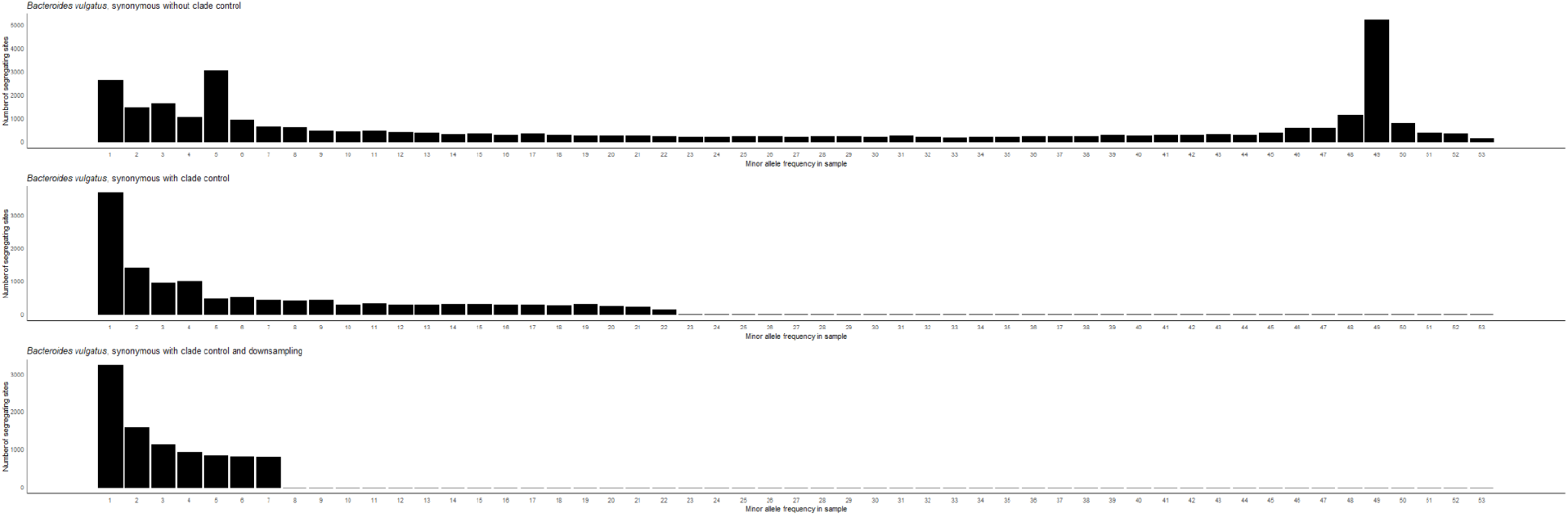
Effects of control and downsampling on the site-frequency spectra. https://github.com/garudlab/microbiome_demography_manuscript/blob/main/Supplement/Supplemental_Figure_2.png Synonymous folded empirical site-frequency spectra of *Bacteroides vulgatus* when (**A**) including lineages from all clades, (**B**) when including lineages only from the largest clade and excluding any closely related lineages, and (**C**) when the samples from (**B**) are downsampled to 14 individuals . Each additional modification to the SFS produces a progressively smoother and more monotonic relationship amongst bins.

**Figure S3:** SFSs and likelihood surfaces for demographic and DFE inference with core genes (see attachment) https://github.com/garudlab/microbiome_demography_manuscript/blob/main/Supplement/Supplemental_Figure_3.png (Left) Site frequency spectra from the empirical data compared to those predicted from the maximum likelihood models for core genes from 27 common commensal gut microbiota. “MLE synonymous” shows the expected SFS produced by the ML demographic parameter estimates. “MLE nonsynonymous” shows the expected SFS produced by the ML demographic and selection parameters from a gamma-distributed DFE. (Right) 2-dimensional likelihood surfaces of population size, ν (in units of N_Anc_) and time since the onset of the most recent demographic event, τ (in units of 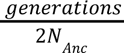). A dashed red line separates the parameter space between contractions and expansions. The maximum-likelihood demographic parameters are shown with an orange dot, and colored intervals denote the decrease in log-likelihood from the MLE. The light cyan regions (LL - 3) correspond to the asymptotic 95% confidence interval, from the chi-squared distribution with 2 degrees of freedom.

**Figure S4:**
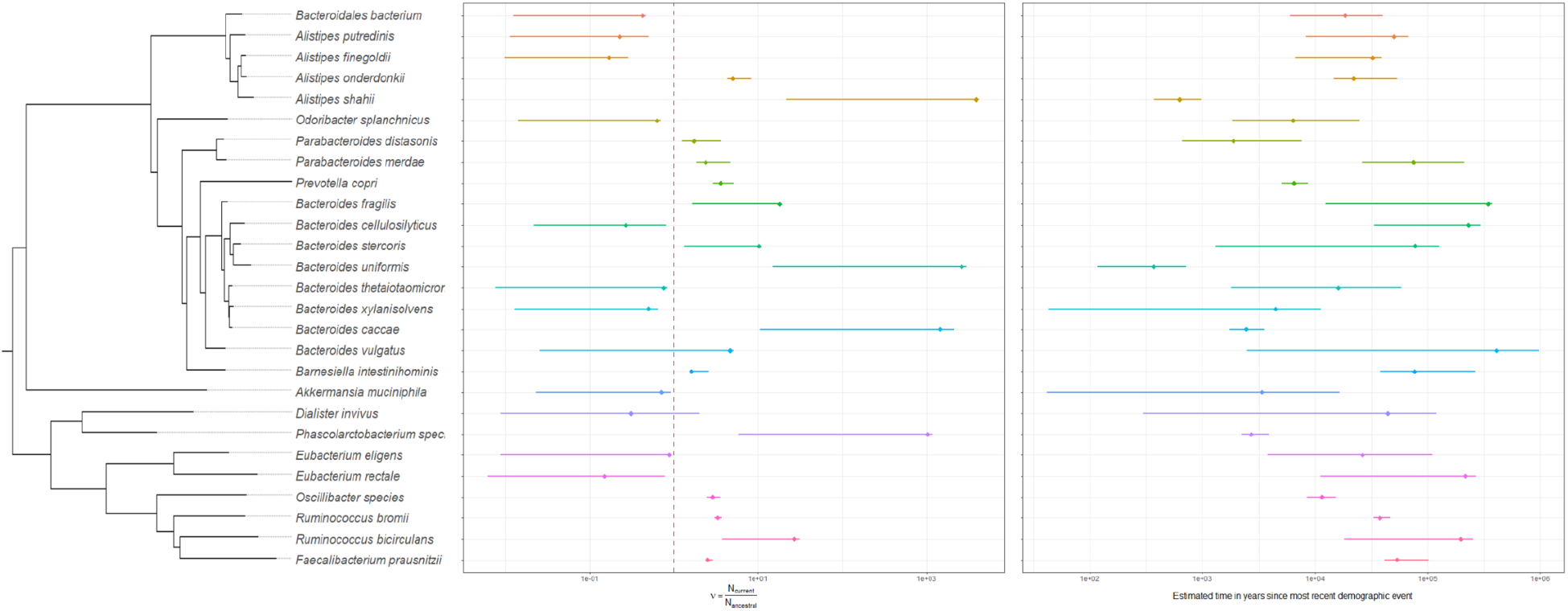
Confidence intervals for ν and τ for demographic inference from core genes. https://github.com/garudlab/microbiome_demography_manuscript/blob/main/Supplement/Supplemental_Figure_4.png Phylogenetically sorted panels of 95% confidence intervals for ν (in units of N_Anc_) (left) and time in years since the most recent demographic event (right). The maximum likelihood parameter is indicated with a diamond. Confidence intervals are color coded by species using the same color scheme as **Figure 3**. For the left plot, a dashed red line separates the parameter space between contractions and expansions.

**Figure S5:**
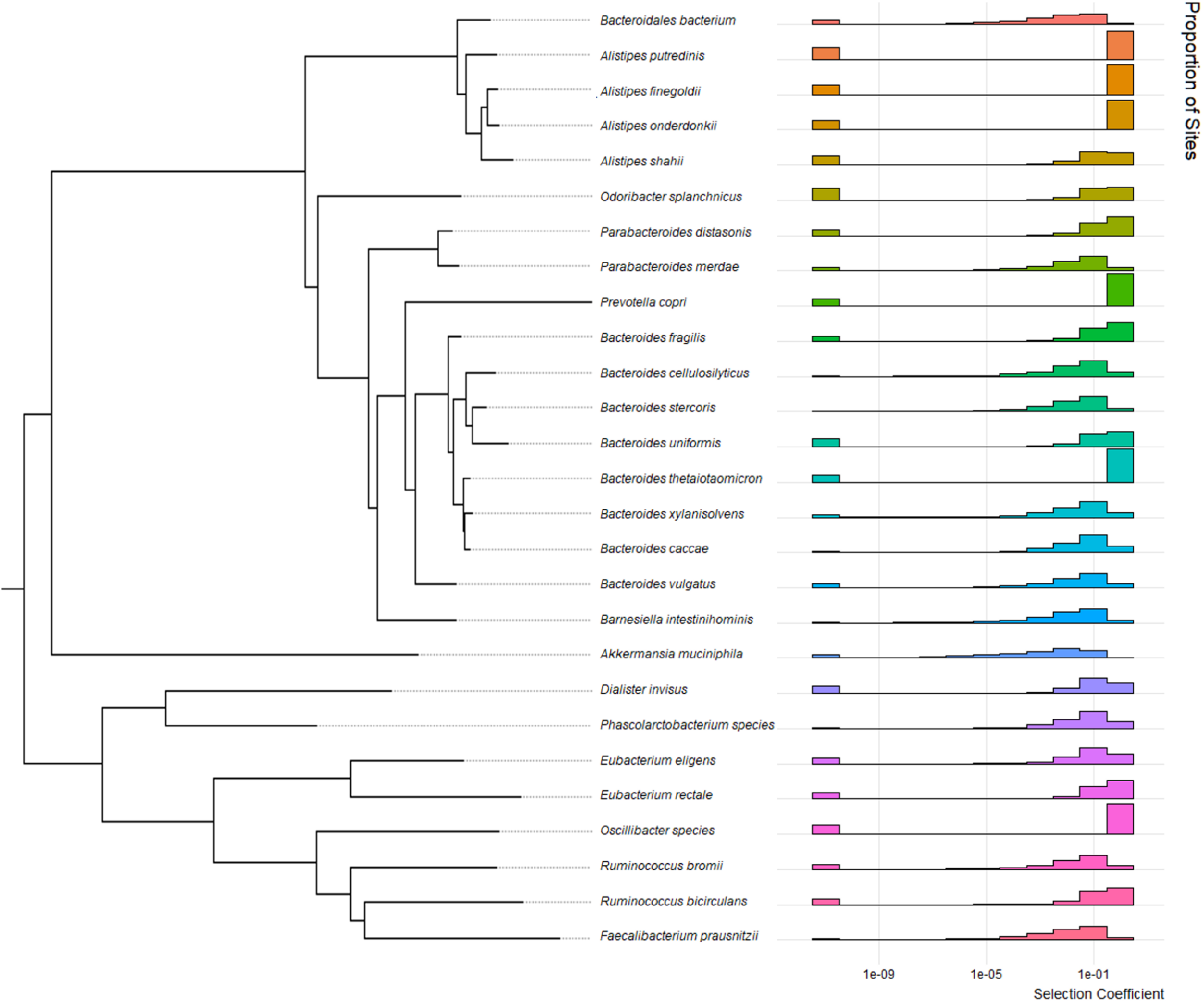
Neu+gamma-distributed DFE inferred from core genes. https://github.com/garudlab/microbiome_demography_manuscript/blob/main/Supplement/Supplemental_Figure_5.png Phylogenetically sorted panel of inferred maximum likelihood distributions of fitness effects per species under a neu+gamma-distribution. With respect to a gamma-distributed DFE, a neu+gamma-distributed DFE is additionally parameterized by *p_neu_,* the proportion of sites assigned to a point mass of neutrality. For each species, the x-axis denotes the log10 scaled discrete bin of selective effect, while the y-axis denotes the proportion of sites which fall into each bin. For ease of visualization, all mutations with selection coefficient, *s*, less than 1 x 10^-12^ are binned in the lowest bin, while all mutations with *s > 0.5* are binned in the highest bin.

**Figure S6:**
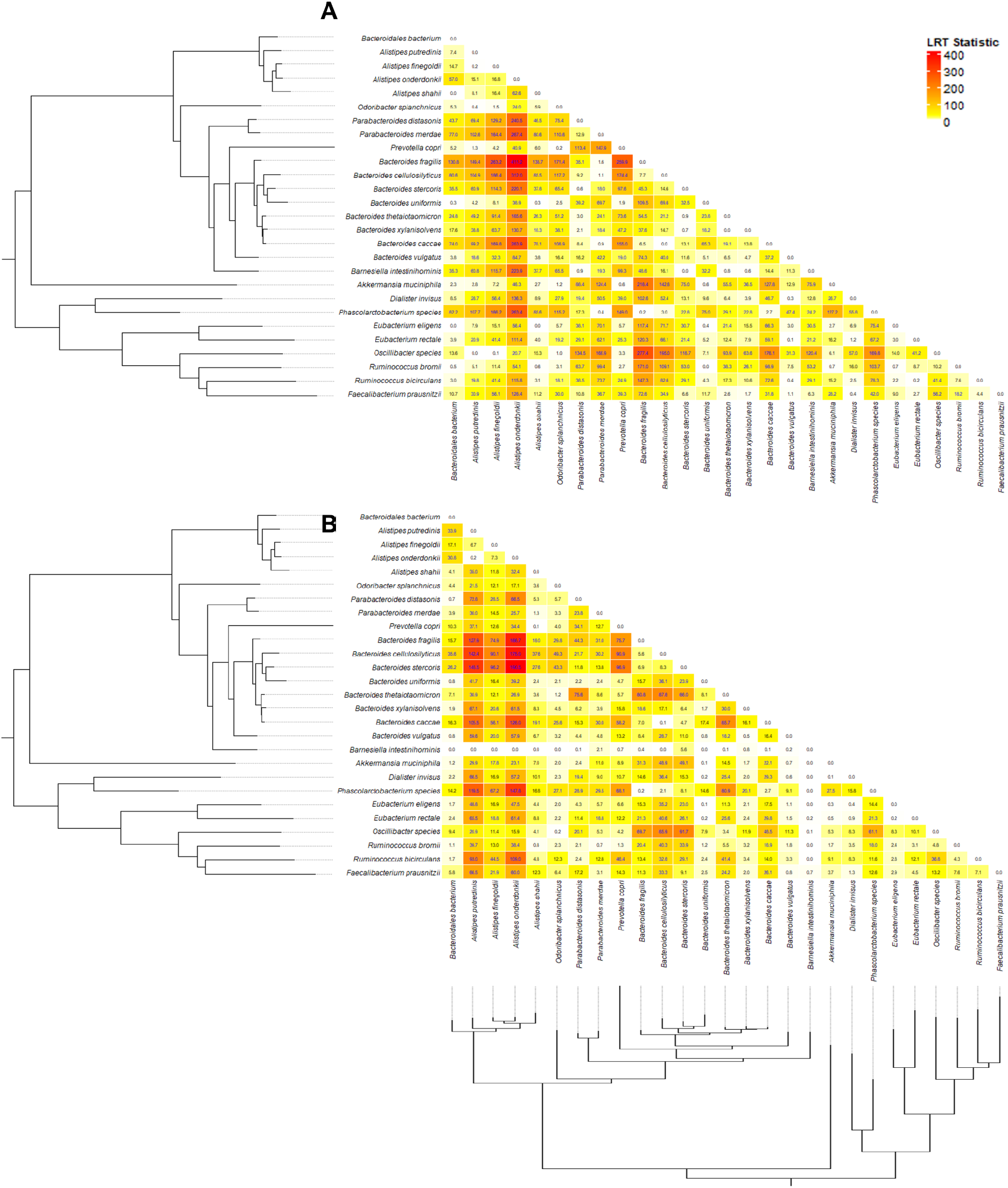
Likelihood ratio test comparing pairs of DFEs in 27 species. https://github.com/garudlab/microbiome_demography_manuscript/blob/main/Supplement/Supplemental_Figure_6.png Each element of the matrix is color coded with a log ratio test statistic comparing a model in which pairs of species have independently inferred DFEs versus a model in which pairs of species have the same DFE. This test was performed in two ways: (**A**) evaluating the selection coefficient, *s,* and (**B**) evaluating the population-scaled selection coefficient 2N_Anc_*s*. A likelihood ratio statistic of about 17.7 corresponds to a 95% confidence interval after Bonferroni correction for 351 tests, assuming that likelihood follows a chi-squared distribution with two degrees of freedom.

**Figure S7:** Empirical SFS of core and accessory genes (see attachment) https://github.com/garudlab/microbiome_demography_manuscript/blob/main/Supplement/Supplemental_Figure_7.png Empirical site frequency spectra for synonymous and nonsynonymous mutations for core genes vs accessory genes. The SFS of accessory genes display a trend of depleted rare variant frequency and increased common variant frequency compared to the SFS of core genes. Additionally, the SFS of accessory genes is much more jagged and less monotonic than the SFS of core genes.

**Figure S8:** SFSs and likelihood surfaces for demographic inference from accessory genes (see attachment) https://github.com/garudlab/microbiome_demography_manuscript/blob/main/Supplement/Supplemental_Figure_8.png (Left) Site frequency spectra from the empirical data compared to those predicted from the maximum likelihood models for accessory genes from 27 common commensal gut microbiota. “MLE synonymous” shows the expected SFS produced by the ML demographic parameter estimates. “MLE nonsynonymous” shows the expected SFS produced by the ML demographic and selection parameters from a gamma-distributed DFE. (Right) 2-dimensional likelihood surfaces of population size, ν (in units of N_Anc_) and time since the onset of the most recent demographic event, τ (in units of 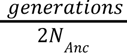) are shown for accessory genes from the same 27 species for which we fit demography. A dashed red line separates the parameter space between contractions and expansions. The maximum-likelihood demographic parameters are shown with an orange dot, and colored intervals denote the decrease in log-likelihood from the MLE. The light cyan regions (LL - 3) correspond to the asymptotic 95% confidence interval, from the chi-squared distribution with 2 degrees of freedom. By visual inspection, demographic inference over accessory genes yields well-fitting models in a minority (n=7) of species.

**Figure S9:**
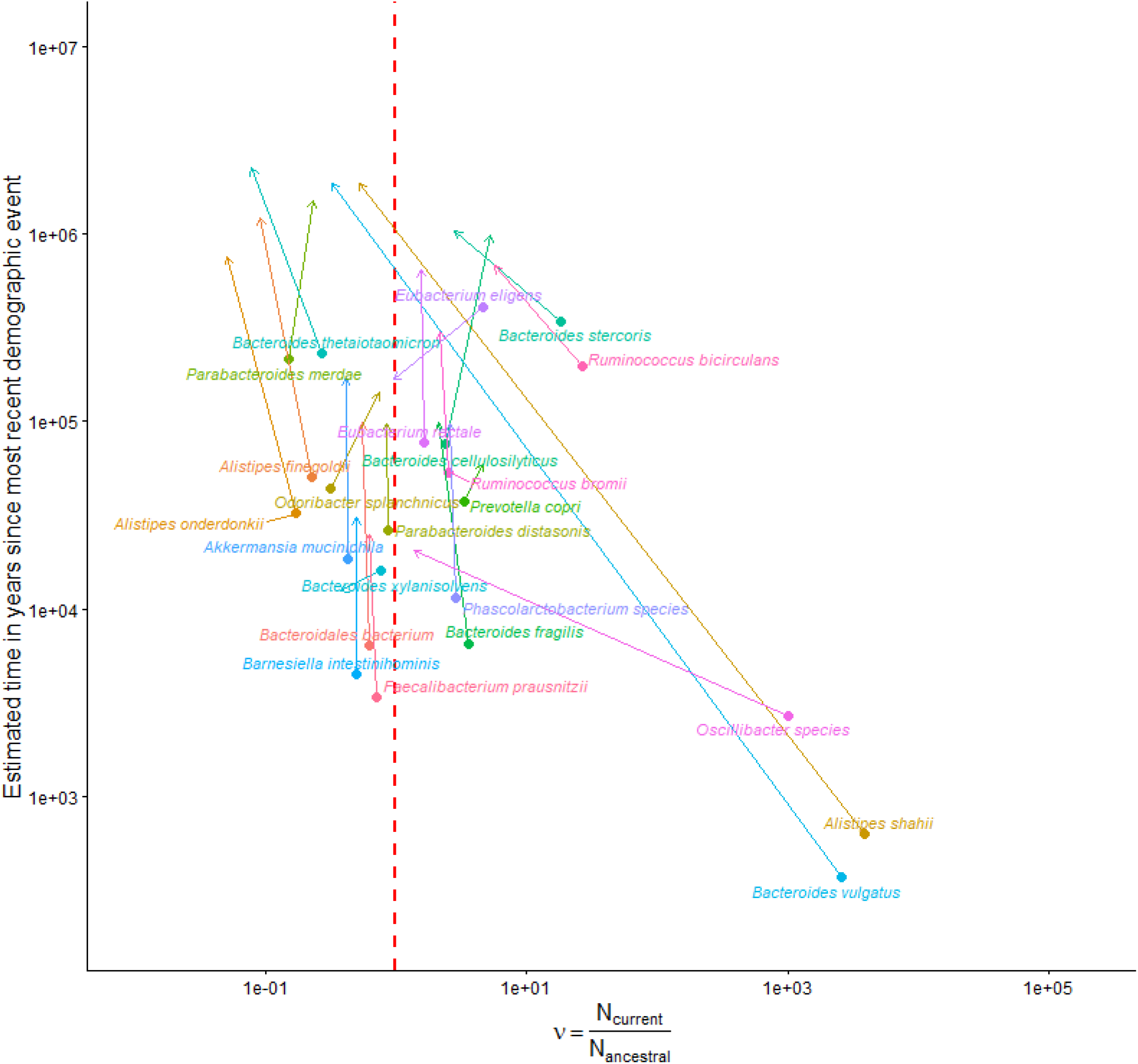
Demography of core genes vs. demography of core + accessory genes. https://github.com/garudlab/microbiome_demography_manuscript/blob/main/Supplement/Supplemental_Figure_9.png A summarizing 2-dimensional scatterplot of ν and approximate time in years for ML demographic parameters for 23 common commensal gut microbiota. A dashed red line separates the parameter space between contractions and expansions. The ML parameters for demographic inference from core genes (circles, labeled by species) is connected by an arrow pointing to the ML parameters for demographic inference from an SFS composed of both core genes and accessory genes (arrow tip). For a majority of species (n=18), we infer a relative decrease in ν when including accessory genes vs. when including only core genes.

**Figure S10:** Phylogenetically ordered analysis of core and accessory gene DFE (see attachment) https://github.com/garudlab/microbiome_demography_manuscript/blob/main/Supplement/Supplemental_Figure_10.png (**A**) Site frequency spectra from the empirical data compared to those predicted from the maximum likelihood models for seven species: *Parabacteroides distasonis, Bacteroides uniformis, Bacteroides thetaiotaomicron, Bacteroides vulgatus, Barnesiella intestinihominis, Eubacterium rectale,* and *Faecalibacterium prausnitzii.* “MLE synonymous” shows the expected SFS produced by the ML demographic parameter estimates. “MLE nonsynonymous” shows the expected SFS produced by the ML demographic and selection parameters from a gamma-distributed DFE. **(B)** Comparison of the inferred gamma-distributed DFE from core genes and accessory genes. For each species, the x-axis denotes the log10 scaled discrete bin of selective effect, while the y-axis denotes the proportion of sites which fall into each bin. Bins are color coded from gray to dark red according to their lethality.

**Figure S11:**
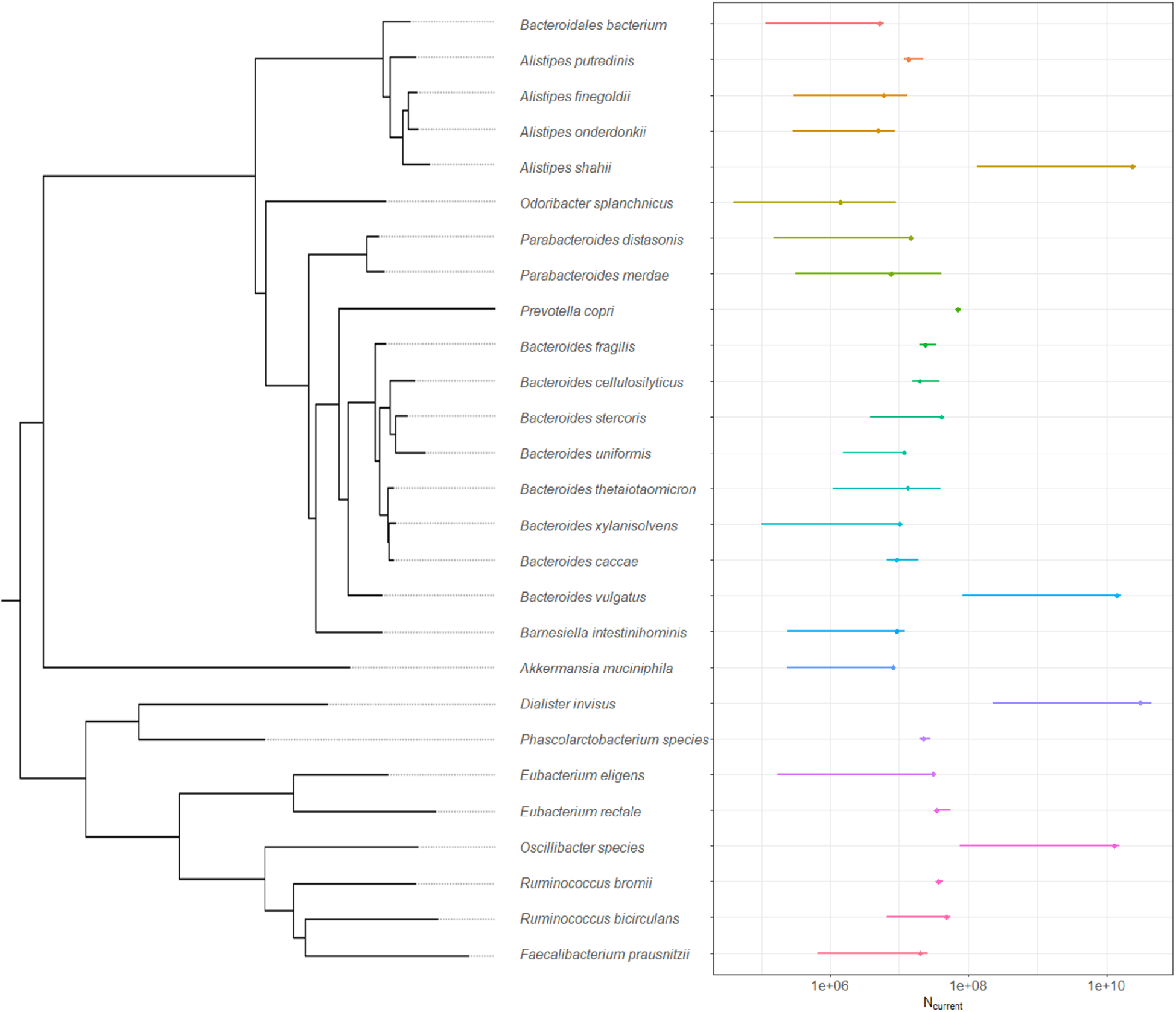
Distribution of N_Curr_ for core genes. https://github.com/garudlab/microbiome_demography_manuscript/blob/main/Supplement/Supplemental_Figure_11.png Phylogenetically sorted panel of 95% confidence intervals N_Curr_. The maximum likelihood parameter is indicated with a diamond. Confidence intervals are color coded by species using the same color scheme as **Figure 3**.

**Figure S12:**
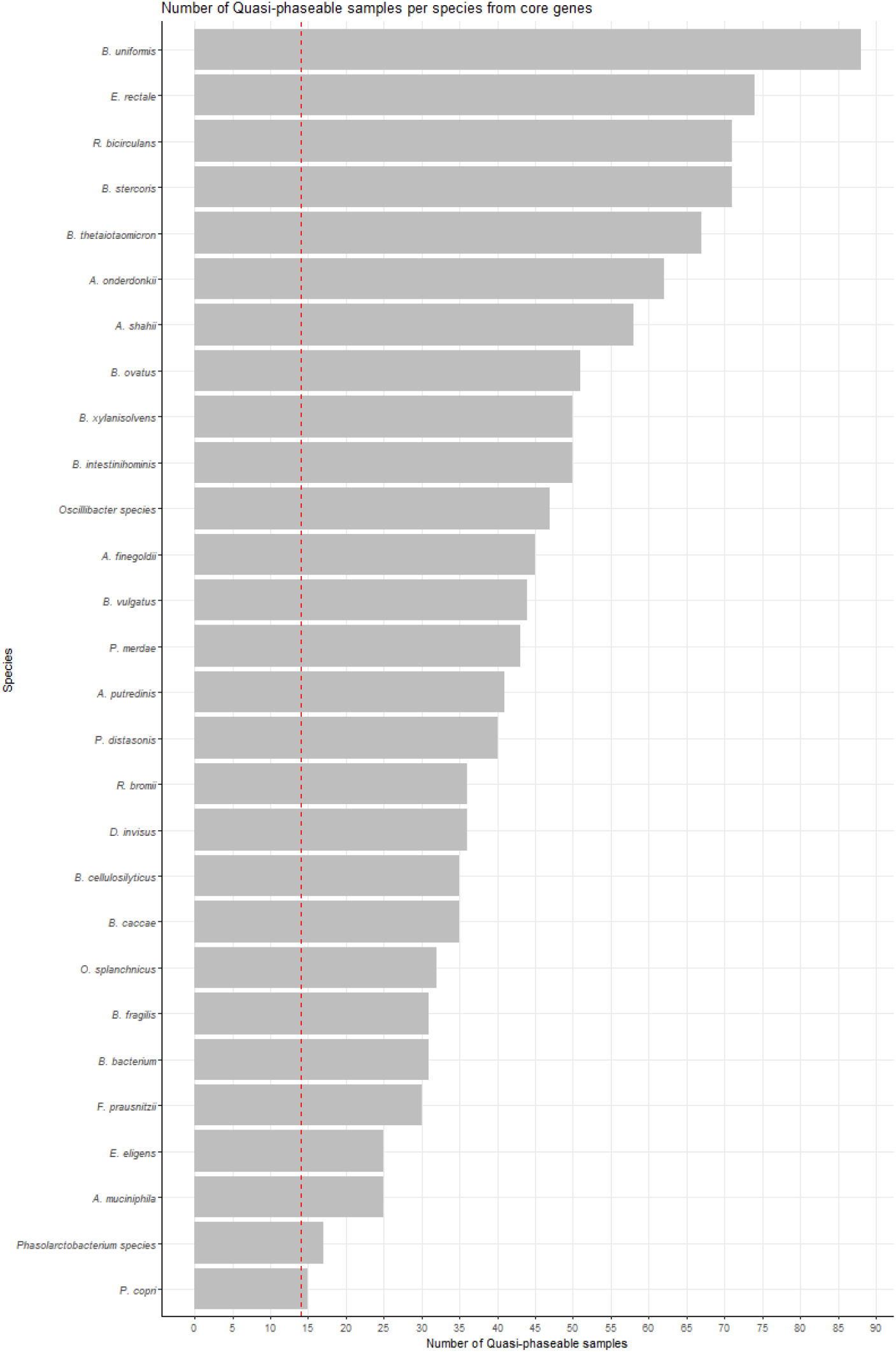
Number of quasi-phaseable samples in top-level clade of core genes per species. https://github.com/garudlab/microbiome_demography_manuscript/blob/main/Supplement/Supplemental_Figure_12.png Species organized in descending order of number of quasi-phaseable samples found from the top-level clade of core genes after removing closely related lineages. A dashed red line indicates the minimum threshold of 14 samples required for use in this analysis.

**Figure S13:**
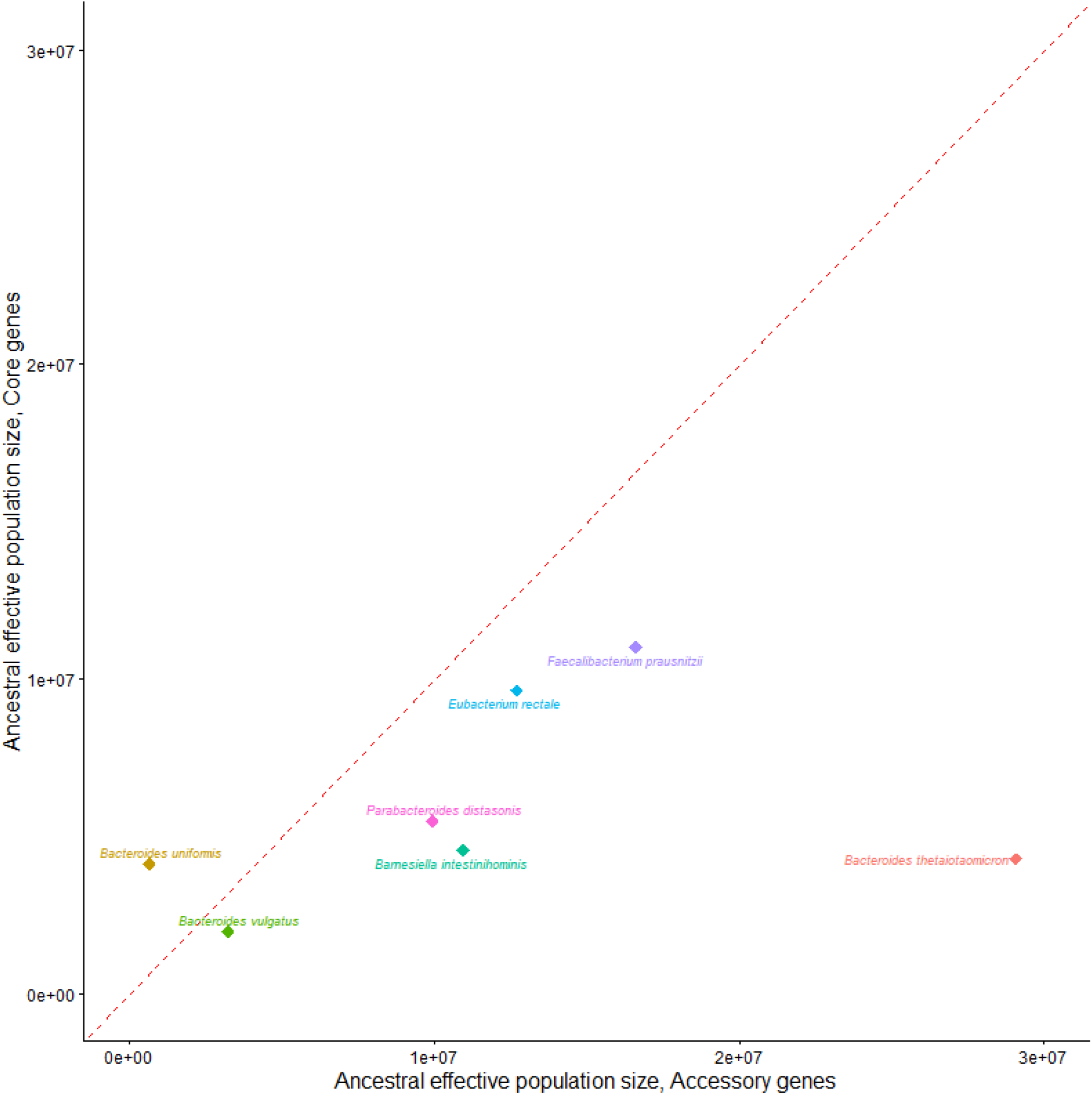
Comparison of N_Anc_ between core and accessory genes. https://github.com/garudlab/microbiome_demography_manuscript/blob/main/Supplement/Supplemental_Figure_13.png 2-dimensional scatterplot comparing the estimated ancestral effective population size of accessory genes vs. core genes. Points are color-coded and labeled by species. A dashed red line indicates a slope of 1.

