## Supplementary figures and images for "Inference of the demographic histories and selective effects of human gut commensal microbiota over the course of human history"

### Supplemental_Figure_1.png

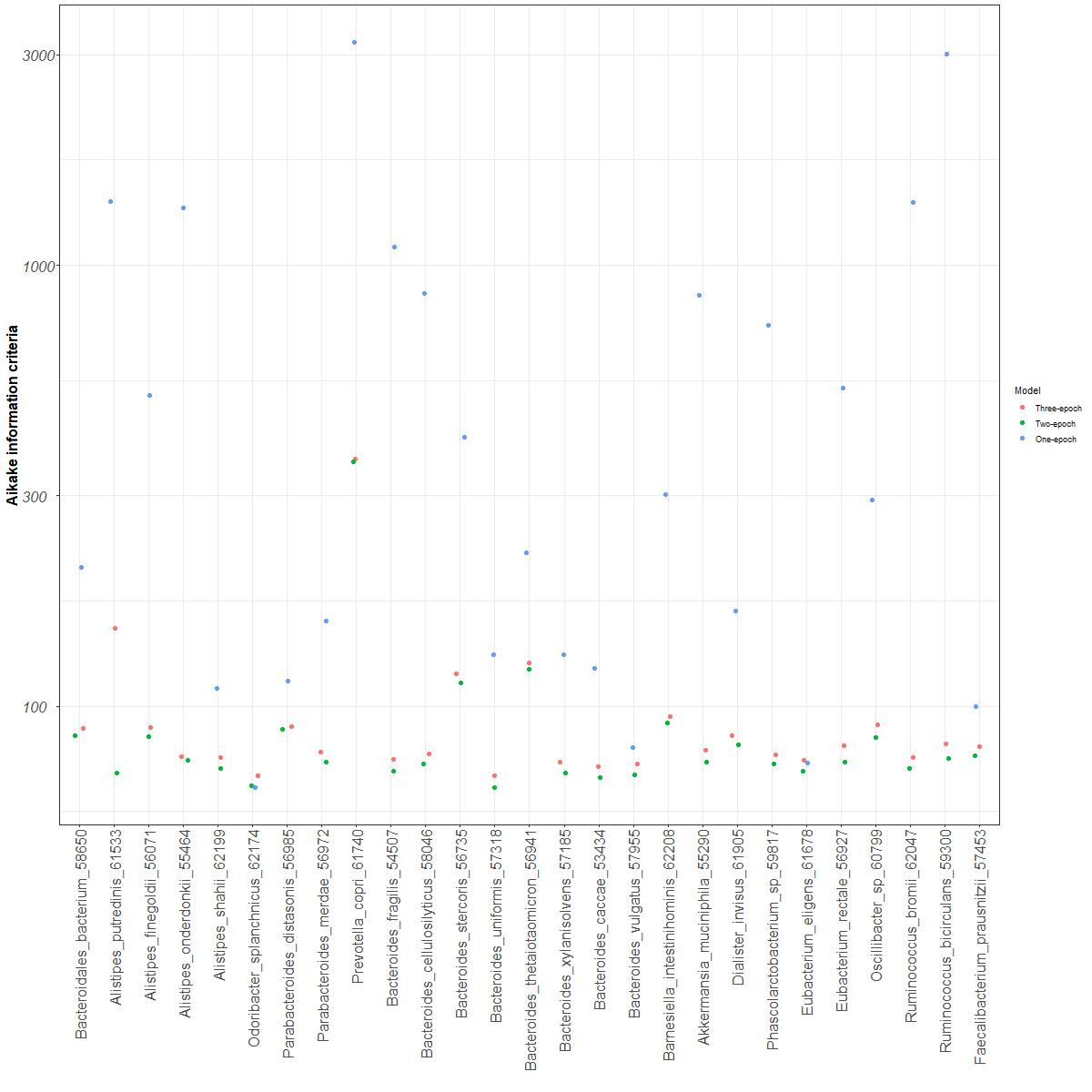

### Supplemental_Figure_2.png

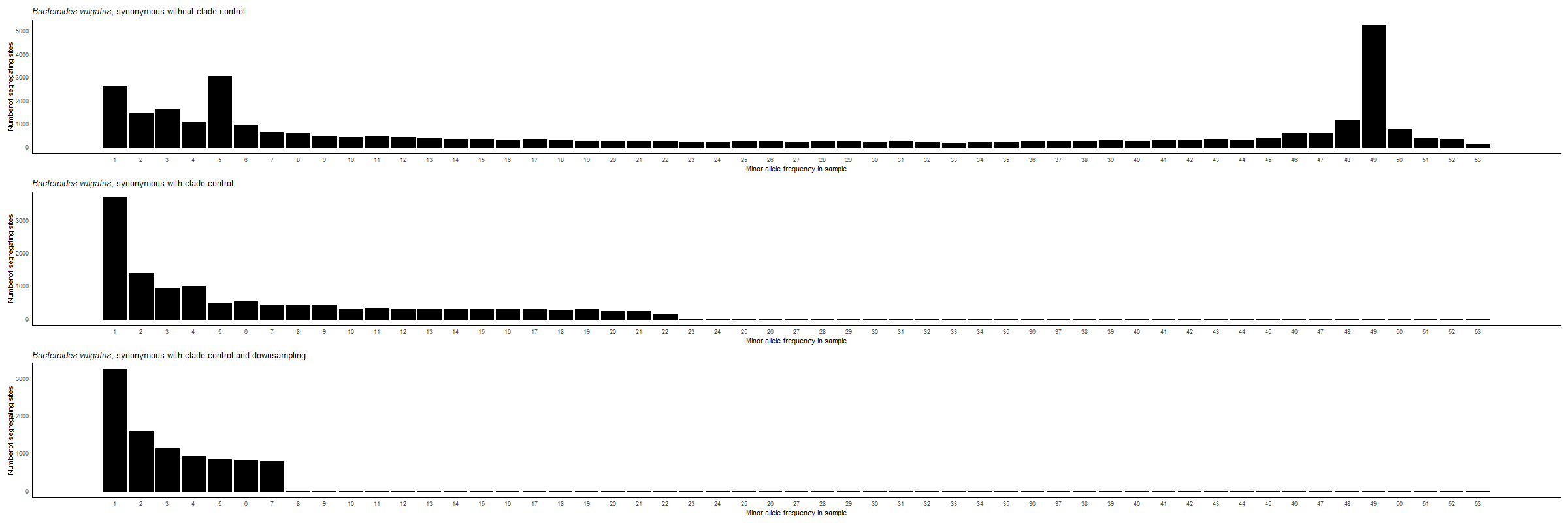

### Supplemental_Figure_3.png

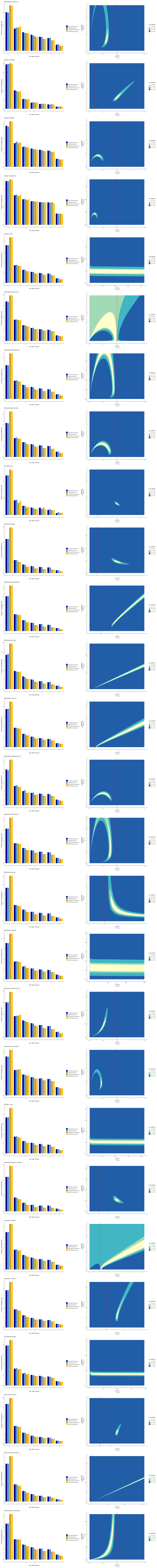

### Supplemental_Figure_4.png

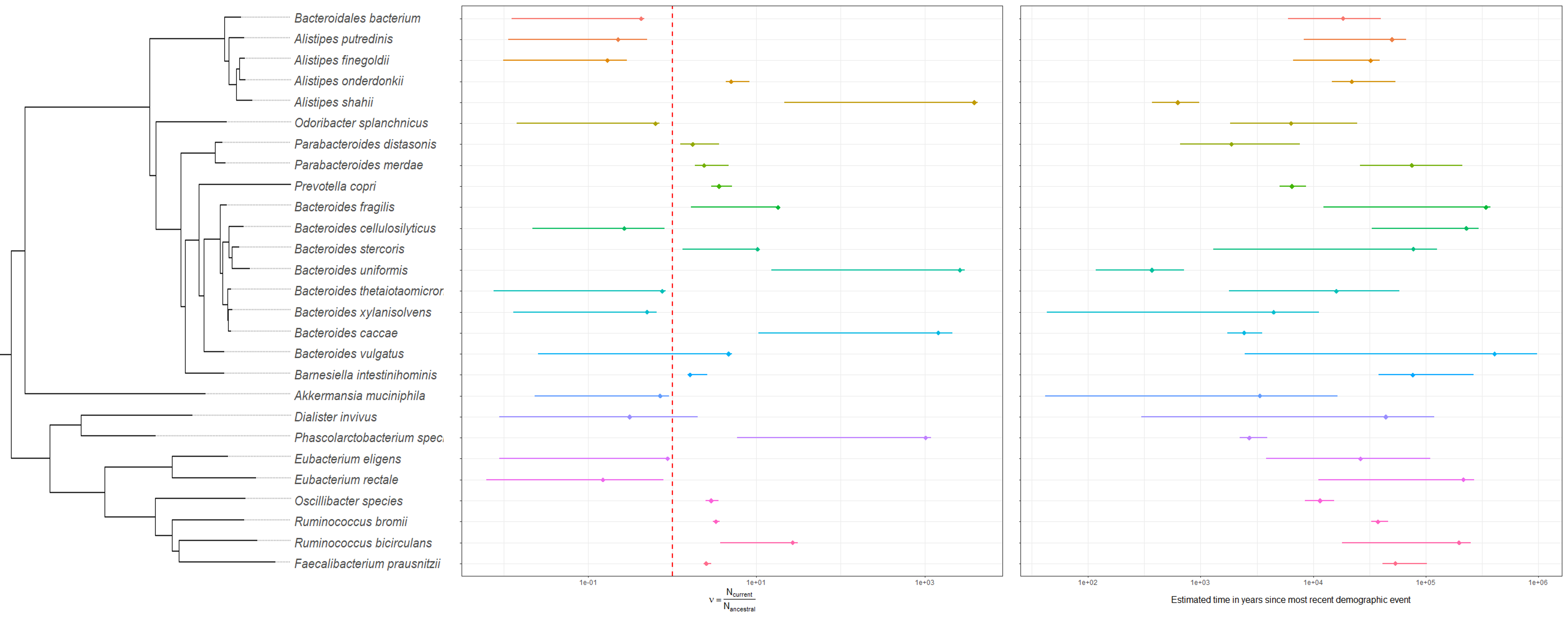

### Supplemental_Figure_5.png

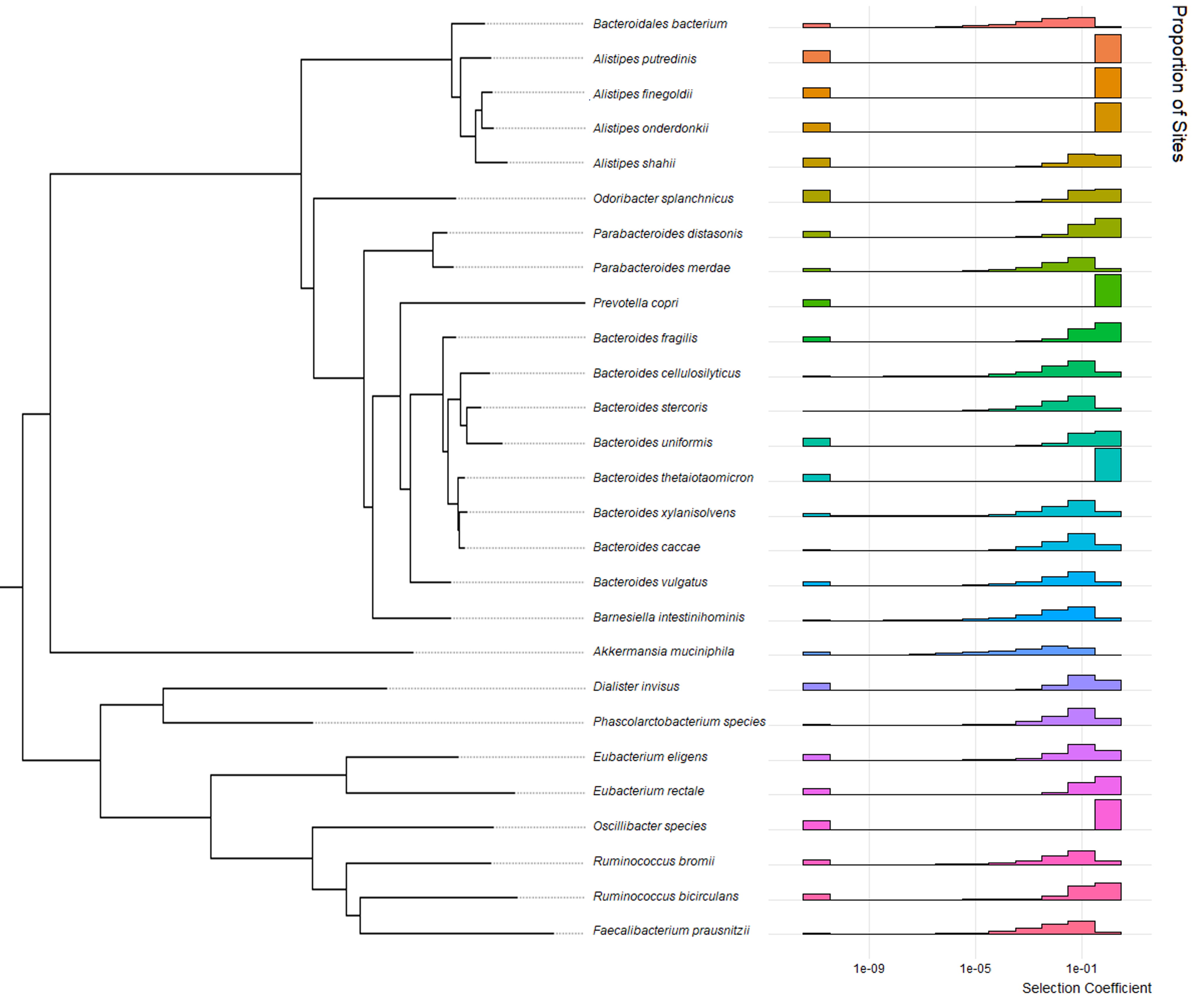

### Supplemental_Figure_6.png

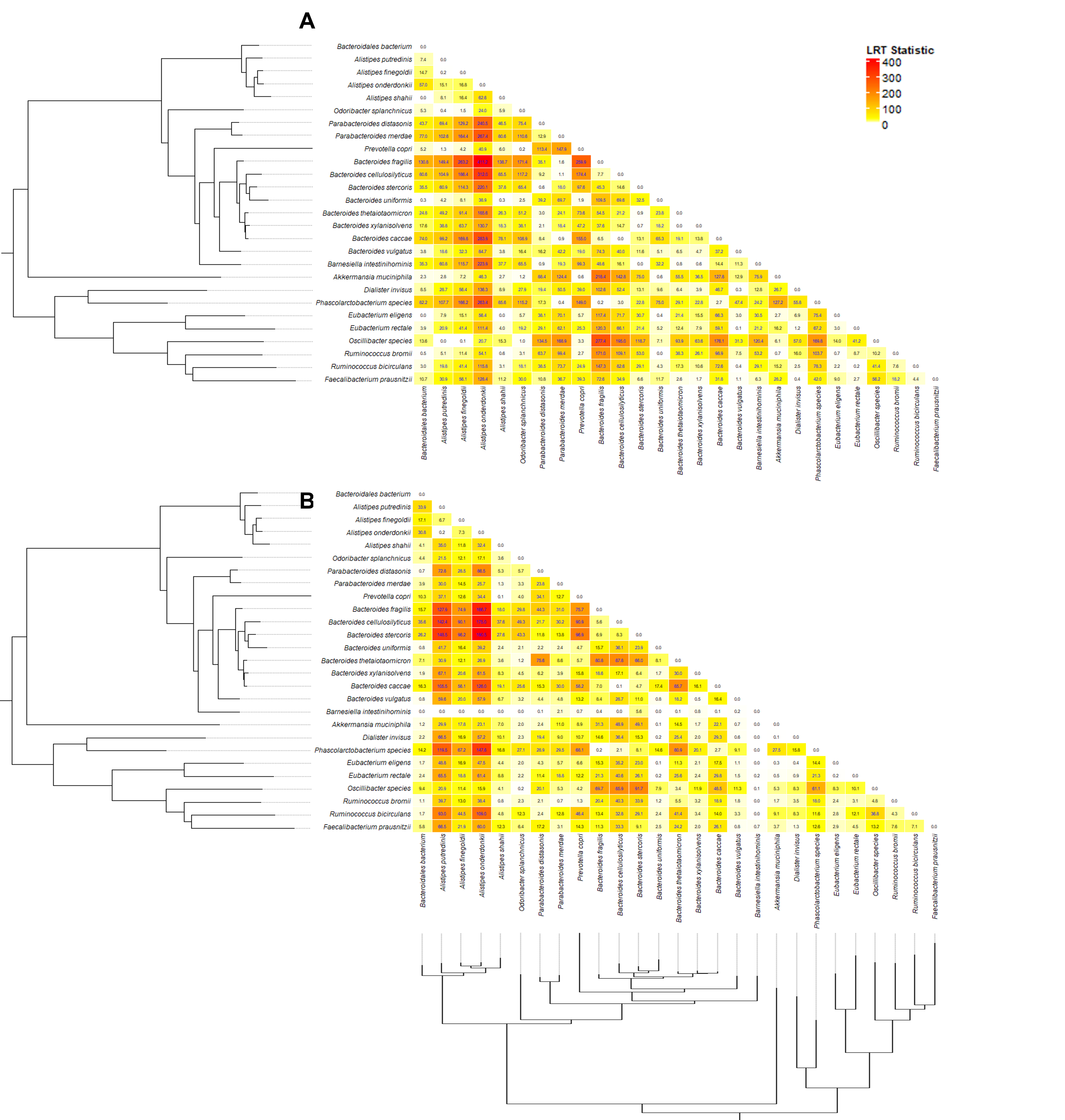

### Supplemental_Figure_7.png

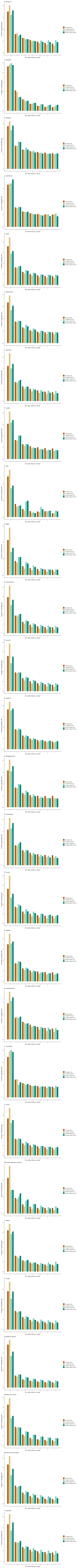

### Supplemental_Figure_8.png

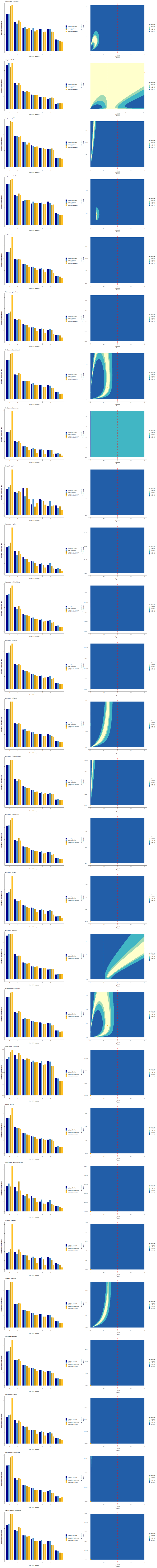

### Supplemental_Figure_9.png

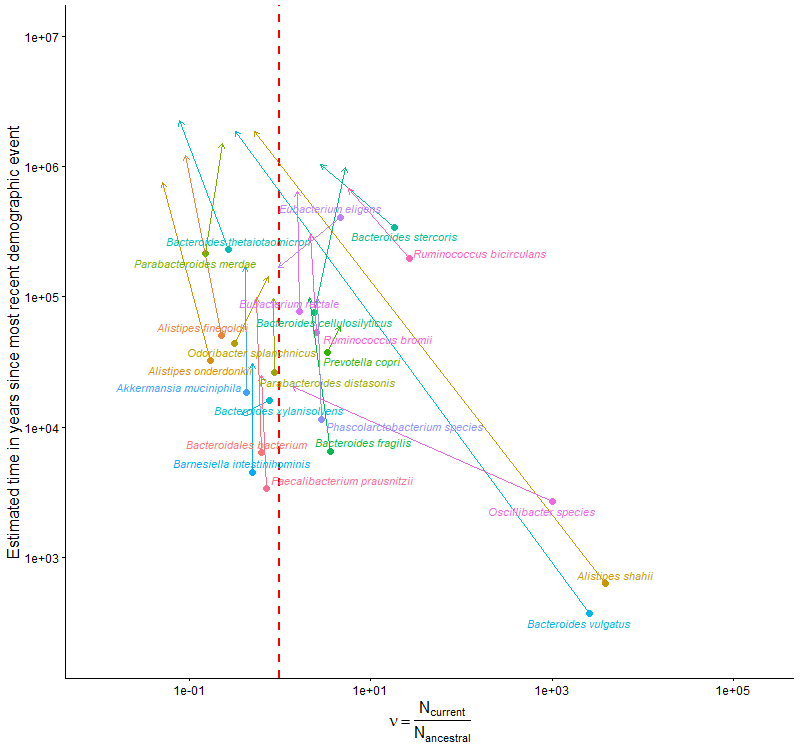

### Supplemental_Figure_10.png

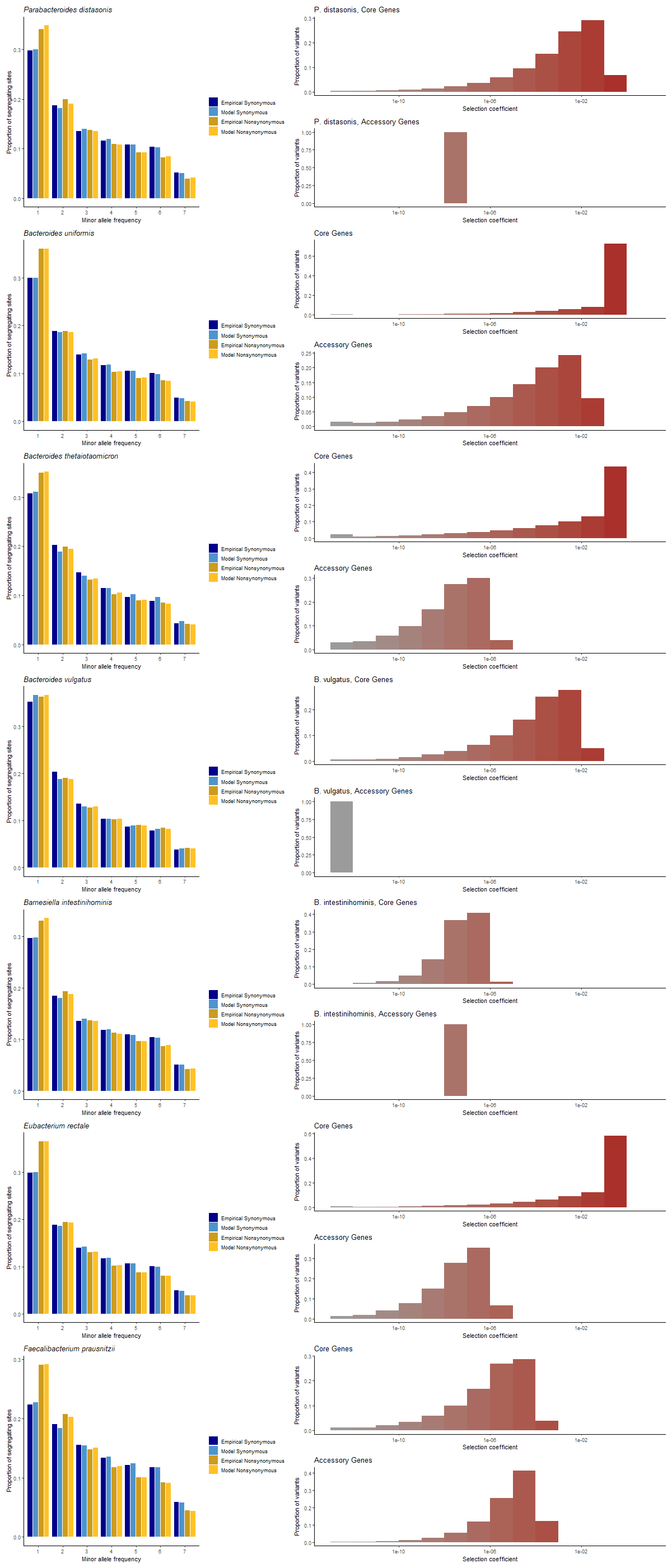

### Supplemental_Figure_11.png

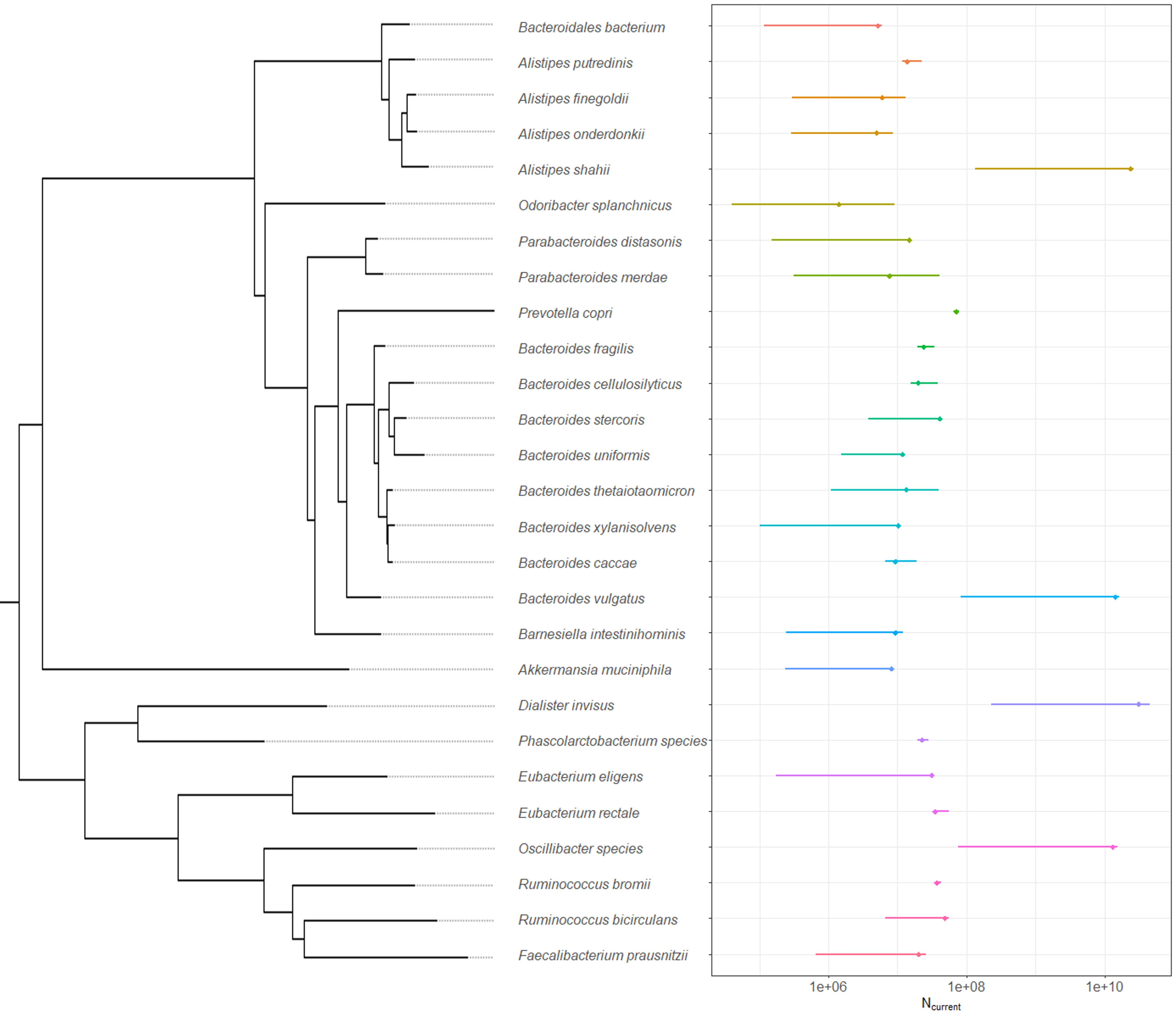

### Supplemental_Figure_12.png

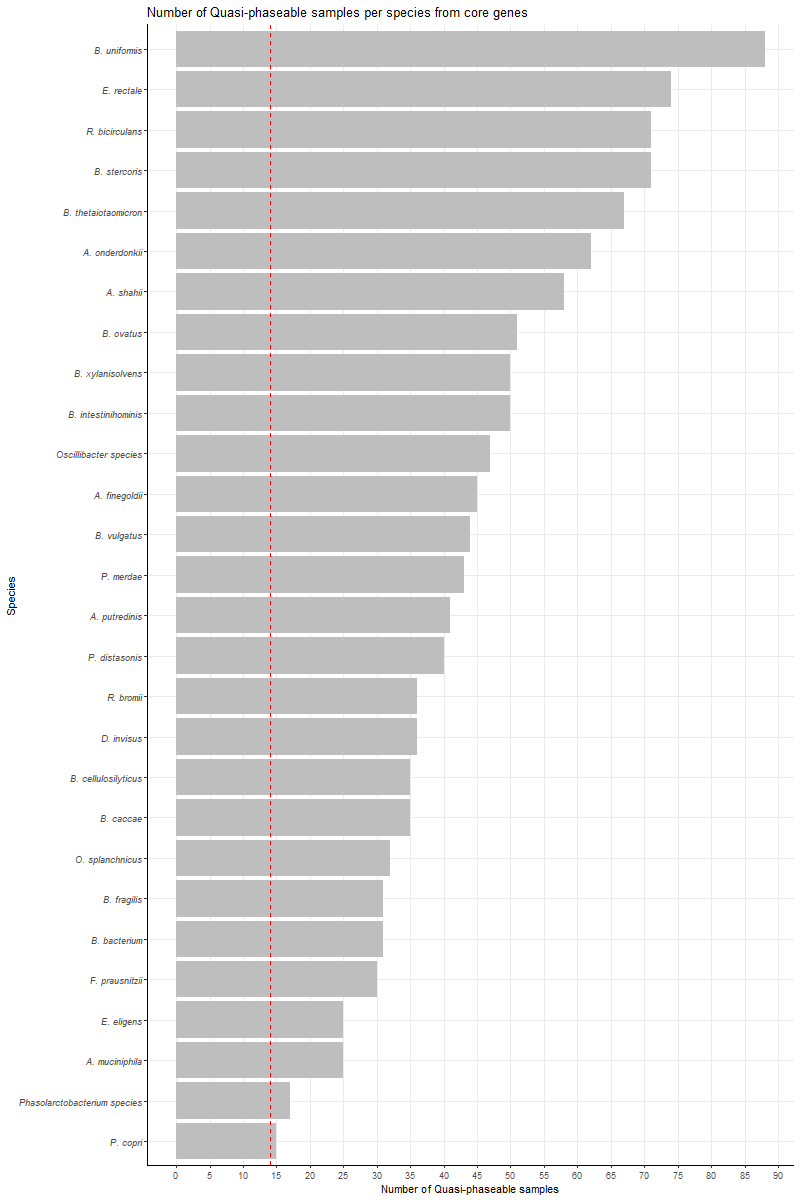

### Supplemental_Figure_13.png

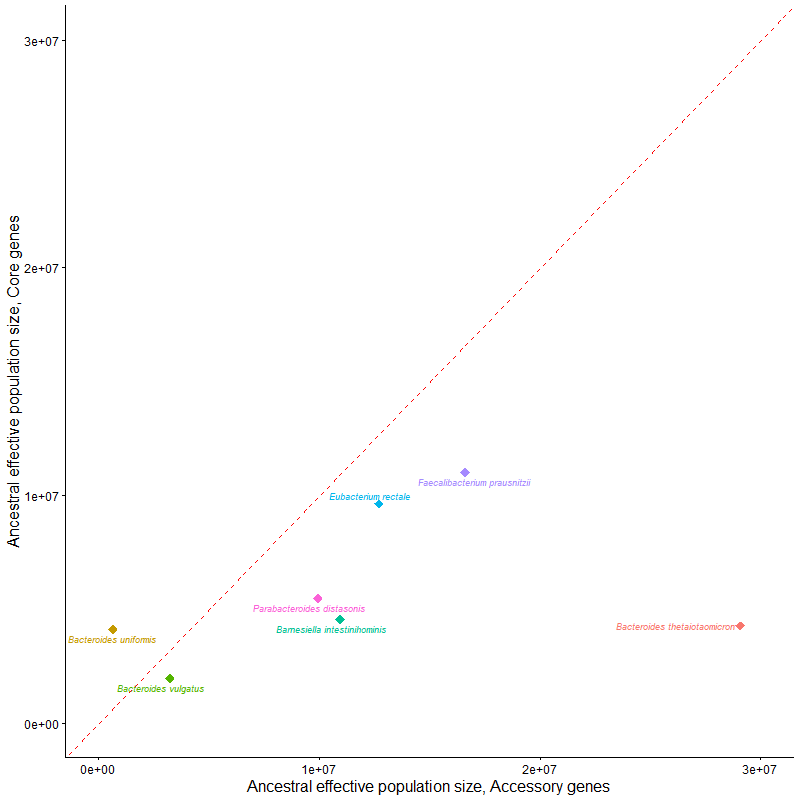
